# HIV Care Prioritization using Phylogenetic Branch Length

**DOI:** 10.1101/2019.12.20.885202

**Authors:** Niema Moshiri, Davey M. Smith, Siavash Mirarab

## Abstract

In HIV epidemics, the structure of the transmission network can be dictated by just a few individuals. Public health intervention, such as ensuring people living with HIV adhere to antiretroviral therapy (ART) and are continually virally-suppressed, can help control the spread of the virus. However, such intervention requires utilizing the limited public health resource allocations. As a result, the ability to determine which individuals are most at-risk of transmitting HIV could allow public health officials to focus their limited resources on these individuals. Molecular epidemiology suggests an approach: prioritizing people living with HIV based on patterns of transmission inferred from their sampled viral sequences. In this paper, we introduce ProACT (**Pr**i**o**ritization using **A**n**C**es**T**ral edge lengths), a phylogenetic approach for prioritizing individuals living with HIV. ProACT uses a simple idea: ordering individuals by their terminal branch length in the phylogeny of their virus. In simulations and also on a dataset of HIV-1 subtype B *pol* sequences obtained in San Diego, we show that this simple strategy improves the effectiveness of prioritization compared to state-of-the-art methods that rely on monitoring the growth of transmission clusters defined based on genetic distance.

---

The transmission of Human Immunodeficiency Virus (HIV) resembles scale-free networks (Wertheim et al., 2014), in which the majority of the structure of the network is dictated by just a few individuals, a phenomenon likely resulting from the scale-free properties of sexual contacts and injection drug use along which HIV is transmitted (Little et al., 2014; Schneeberger et al., 2004). As a result, public health intervention may be more effective when targeted at people living with HIV (PLWH for short) who are more likely to grow the transmission network. However, the best method to target individuals for specific interventions remains an open question, and the best strategy will likely depend on the specific intervention planned.

A potential form of intervention aiming to reduce future transmissions is to target PLWHs. Antiretroviral therapy (ART) is an effective treatment of HIV that suppresses the HIV virus in the majority of cases, stops the progression of the disease, and prevents onward transmission to an uninfected sexual partner, provided the PLWH continuously adheres to the treatment (Cohen et al., 2011). In most advanced health care systems, ART is made available routinely to newly diagnosed patients, but several opportunities for further intervention remains available. Most importantly, not every diagnosed person initiates ART and not all cases of ART initiation lead to a sustained suppression of the virus through time. PLWHs who start ART but fail to sustain it or who are otherwise unsuppressed can still infect others. Thus, a possible intervention is to use public health resources to help known PLWHs stay on ART and to remain continually suppressed (Poon et al., 2016). Such interventions require allocation of clinical staff who would follow up with patients to provide them further assistance in adherence sustenance of ART. They health system can also provide increased testing to these individuals to ensure suppression. A second family of interventions involves targeting HIV negative individuals connected to high priority PLWHs. The health system can use partner tracing (Gotz et al., 2014) to identify the sexual partners of high-priority PLWHs (as best as possible), test these high risk individuals, and offer them either treatment (for positives) or prevention through PrEP (for negatives). Finally, if the priority status of individuals shows any association with specific geographical or demographic groups (beyond known associations), the public health system can design strategies for further outreach, testing, and PrEP administration for the impacted groups.

All three types of intervention are costly and cannot be undertaken for every known PLWH or groups. If diagnosed people at risk of not being suppressed could be predicted accurately, the public health system could focus their limited resources on these individuals, Thus, a natural question surfaces: which individuals are most at-risk of transmitting HIV? However, predicting tendency for future transmissions is difficult and can also be problematic if undertaken primarily based on demographic or behavioral traits.

Molecular epidemics suggest an alternative method: prioritizing PLWHs for intervention solely based on patterns of transmission inferred from HIV sequence data (Bbosa et al., 2019; Villandré et al., 2019; Oster et al., 2018; Ragonnet-Cronin et al., 2019; Wertheim et al., 2018, 2011, 2014; Smith et al., 2009). The inference of transmission networks using phylogenetic or distance-based methods has been the subject of much research (e.g. Leitner and Romero-Severson, 2018; Kosakovsky Pond et al., 2018; Ragonnet-Cronin et al., 2013; Prosperi et al., 2011). However, in this work, instead of being concerned with inferring exact patterns of transmissions, we ask the following question: given molecular data from a set of *sequenced* PLWHs (“samples” for short), who should be prioritized for further intervention?

Prioritizing care based on molecular epidemics has been studied recently. Wertheim et al. (2018) present a method for prioritizing samples based on performing transmission clustering (i.e., grouping individuals with low viral genetic distance into *transmission clusters*) and ordering clusters by growth rate. On a large dataset from New York, they show that the approach is able to predict individuals who have relatively larger numbers of transmission links in the near future. Moshiri et al. (2018) have studied the same question in simulations and have shown that monitoring cluster growth can be used for predicting future transmissions substantially better than a random guess, whether clusters are defined using genetic distances or using phylogenetic methods. Most recently, Balaban et al. (2019) showed in simulations that using a cluster-monitoring approach similar to that of Wertheim et al. (2018) but defining clusters using a min-cut optimization problem gives a small but consistent improvement over defining clusters using genetic distances.

In this paper, we introduce a new method for ordering samples based on their phylogenetic relationships. Instead of relying on clustering individuals and then ordering clusters based on their growth, we seek to order individuals without clustering and without reliance on parametric models. Instead, we seek to simply exploits patterns in the phylogeny, and in particular, in branch lengths.

## Materials and Methods

ProACT (**Pr**i**o**ritization using **A**n**C**es**T**ral edge lengths) takes as input the inferred phylogenetic relationships between sampled HIV viruses (e.g. from the *pol* region), rooted using an outgroup or clock-based methods (e.g. midpoint or MinVar-root, Mai et al. (2017)). ProACT simply orders samples in order of incident branch length of their associated virus, and it breaks ties based on incident branch lengths of parent nodes, then those of grandparent nodes, etc. We first motivate the approach and then present a formal definition of the method.

We note that ProACT is motivated and tested in a context similar to the present day health care systems that enjoy enough resources to provide ART to all (or at least most) diagnosed individuals. Thus, each sample can be assumed to be given ART at a time close to when their HIV is sequenced, but they may fail to be suppressed for the remainder of their life. These conditions describe the common practice of care in many advanced and (increasingly) developing countries.

### Motivating the Approach

We start with the observation that, in simulations (described in detail below), when a phylogeny is inferred from sequences obtained at a given time point in an epidemic, the more a node transmits, the shorter its incident branch length tends to be (Figs. 1d–e and S2). Using the Kendall’s Tau-b test (Kendall, 1938), in a ten-year epidemic simulation (details described below), we found a statistically significant anticorrelation between the incident branch lengths of individuals sampled within the first 9 years of the epidemic and the number of individuals they infected over the final year of the epidemic. This held for true *(τ_B_* = −0.0431, *p* ≪ 10^-10^) and inferred *(τ_B_* = −0.0354, *p* ≪ 10^-10^) phylogenetic trees. Though not obvious, this observation can be explained by the constraints placed upon the viral phylogeny by the transmission history (Fig. 1a-c).

**Fig. 1.**
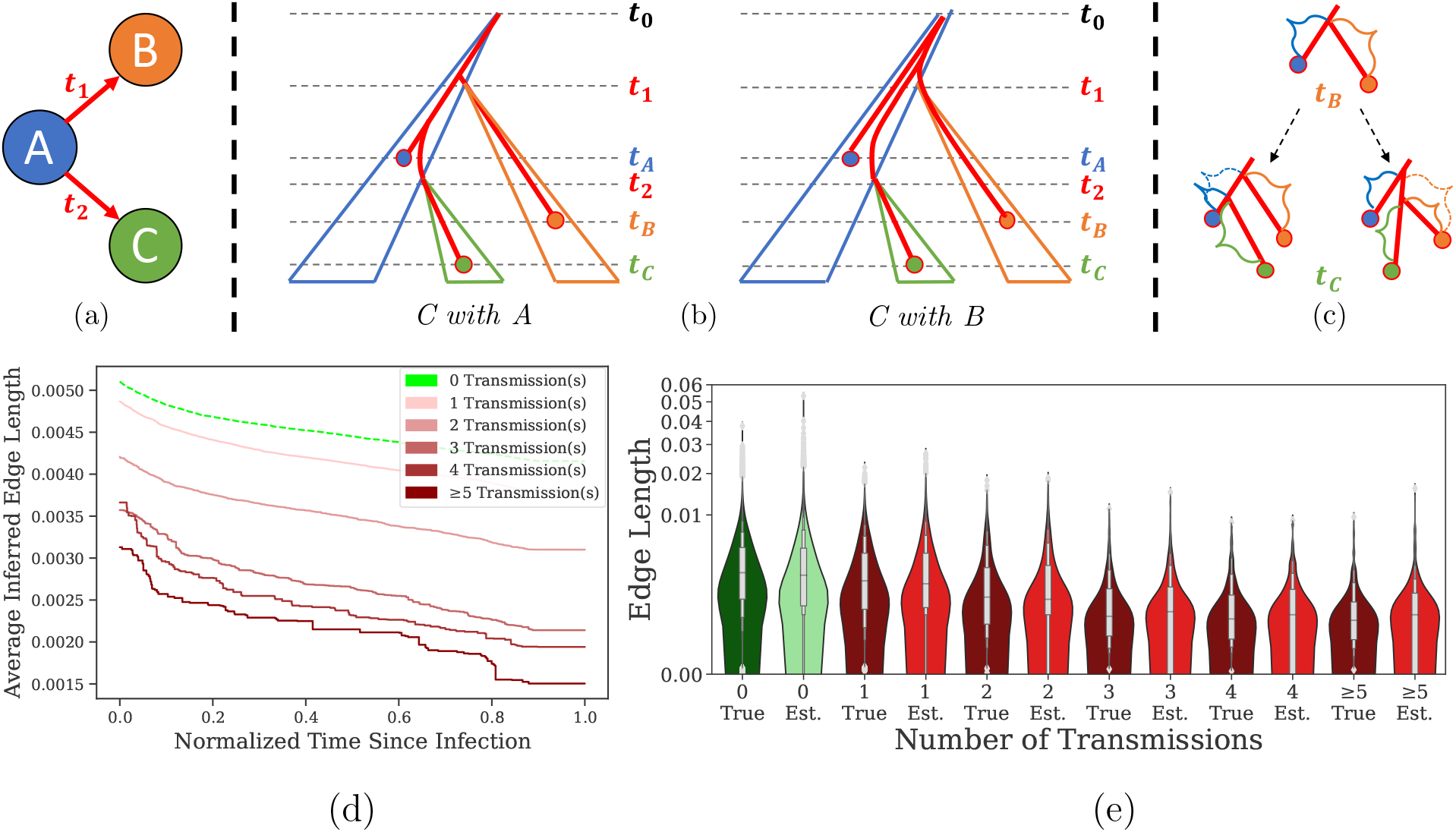
The effect of new transmissions on incident branch lengths. (a) Individual *A* transmits to individual *B* and *C* at times at *t*_1_ and *t*_2_, respectively. (b) Viral samples are obtained from individuals *A, B*, and *C* at times *t_A_, t_B_*, and *t_C_*. The viral phylogeny of samples is constrained by each transmission event’s bottleneck, and the most likely phylogeny matches the transmission history (Left), but in the less likely deeper coalescence, it may not match (Right). (c) Moving from the phylogeny observed at time *t_B_* to the phylogeny at time *t_C_*, the branch length incident to individual A shortens upon the addition of individual C in the likely event that the coalescence of the lineage from *C* with the lineage from *A* is more recent than its coalescence with the lineage from *B* (Left), or the branch length incident to individual A remains constant in the event of a less likely deeper coalescence (Right). Regardless, the length of the branch incident to individual A never increases. In simulation, we can observe this trend: as time progresses, the incident branch length of each individual tends to decrease, both in true (Fig. S1) and inferred (d) phylogenies, and as the number of transmissions from a given individual increases, the distribution of incident edge length tends to decrease, both in true and inferred phylogenies, labeled “True” and “Est.,” respectively (e).

In the context of HIV epidemiology in many advanced countries, samples are typically sequenced upon beginning Antiretroviral Therapy (ART). Let’s assume for simplicity that every individual in the given dataset has at some point initiated ART, meaning future transmissions by individuals in the dataset must happen only if the source stops ART or is otherwise unsuppressed. Given a viral phylogeny containing all known samples, if, in the future, individual *u* in the dataset transmits to individual *v*, there are two possible scenarios regarding the placement of the leaf corresponding to v in the existing (true) phylogeny: (1) *v* is placed on the edge incident to *u*, so the edge incident to *u* will shorten, or (2) *v* is not placed on the edge incident to *u*, so the edge incident to *u* will remain the same length. Although Scenario 2 is possible, Scenario 1 is far more likely (Romero-Severson et al., 2016), and note that the terminal branch lengths do not increase in either scenario. Thus, as time goes by, the terminal branch can only shorten or stay fixed, and it will most often shorten because of new transmissions by the sample associated with that terminal branch. This pattern, easily observed in simulations (Fig. 1d), leads to shorter branches for samples who have transmitted recently.

Note that samples who transmit are unsuppressed. The first time they infect others, their terminal branch length is likely to decrease, and further transmissions further decrease their terminal branch lengths (Fig. 1d). Thus, one expects nodes with smaller incident branch length to be more likely to have transmitted since their sampling time. Moreover, they are also likely to transmit in the near future because they are likely not to be suppressed. The higher probability of a lack of suppression makes them a good candidate for intervention.

### Formal Description

ProACT takes as input a *rooted* phylogenetic tree *T* of viral samples. Let *bl*(*u*) denote the incident branch length of node *u*, and assume the incident branch length of the root of *T* is 0. Let *a*(*u*) denote the vector of ancestors of node *u* (including *u*), where *a*(*u*)_1_ is *u, a*(*u*)_2_ is the parent of *u, a*(*u*)_3_ is the grandparent of *u*, etc. Let *r*(*u*) denote the length of the path from node *u* to the root of *T*, i.e., *r*(*u*) = Σ_*v*∈*a*(*u*)_ *bl*(*v*). ProACT sorts the leaves of *T* in ascending order of *bl*(*a*(*u*)_1_), with ties broken by *bl*(*a*(*u*)_2_), then by *bl*(*a*(*u*)_3_), etc. Note that, for two leaves *u* and *v*, |*a*(*u*)| may be less than |*a*(*v*)|, in which case, for all 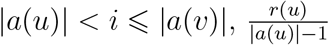 (i.e., average branch length along the path from *u* to the root of *T*) is compared with *bl*(*a*(*v*)_*i*_) instead. If two nodes are equal in all comparisons, if the user provides sample times, the earlier sample time is given higher priority; otherwise, ties are broken arbitrarily. Because sorting is needed, for a tree with *n* leaves, assuming branch lengths are fairly unique, the ProACT algorithm runs in 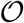(*n* log *n*) time. Scalable methods exist both for the inferring (e.g. Price et al., 2010; Nguyen et al., 2015) and rooting (e.g. Mai et al., 2017) very large trees.

## Results

We evaluate ProACT on simulated and real data.

### Simulation Results

In order to test ProACT’s efficacy, we performed a series of simulation experiments in which we used FAVITES (Moshiri et al., 2018) to generate a sexual contact network, transmission network, viral phylogeny, and viral sequences emulating HIV transmission in San Diego from 2005 to 2014 (Material and Methods). We have simulated nine model conditions (Table 1) by starting from a base model condition and varying the rate of ART initiation (λ_+_), rate of ART termination (λ_−_), and the expected degree of the sexual network (*E_d_*). We subsequently inferred and rooted a phylogeny of all sequences obtained during the first 9 years of the simulation. Then, ProACT was run on the true and inferred full trees and subsampled trees.

**Table 1.**
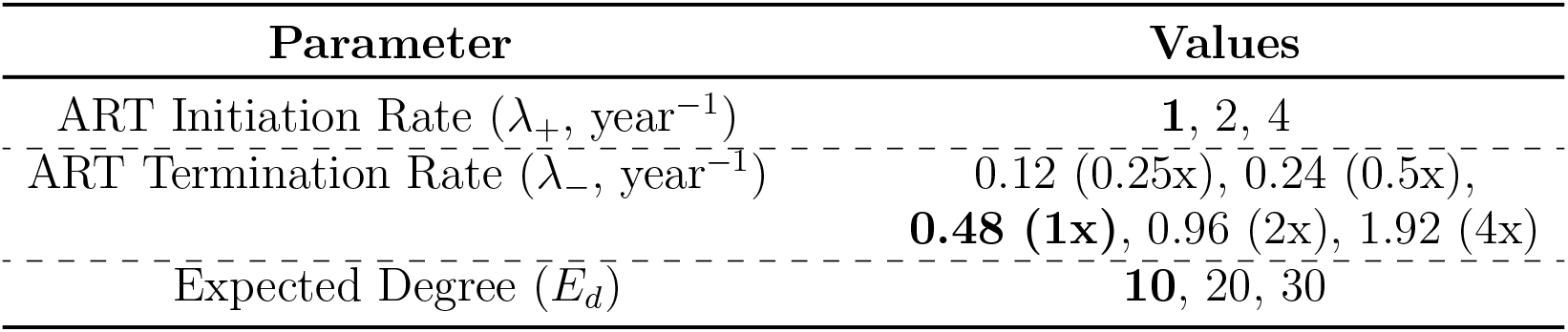
Varied HIV simulation parameters. Values for the base model condition are shown in bold.

To measure the efficacy of a given prioritization, we compute the number of infections caused by each individual during the 10th year of the simulation (our outcome measure). Then, we measure the cumulative moving average (CMA) of the outcome measure by the top samples. The higher the CMA in a prioritization, the higher the number of future transmissions from these top individuals, and thus, the higher the effectiveness of the prioritization. Moreover, sorting individuals by their outcome measure (known to us in simulations) enables us to compute the optimal CMA curve, and the mean number of transmissions gives us the expected value of the CMA for a random prioritization. Across experimental conditions, the maximum and random expectations vary. Thus, to enable proper comparison of effects of prioritization across conditions, we also report an adjusted CMA normalizing above the random prioritization and over the optimal prioritization (see Materials and Methods). For this Adjusted Transmissions/Person metric, 1 indicates the optimal ordering and 0 indicates an ordering that is no better than random (a negative value indicates an ordering that is *worse* than random). Finally, we use Kendall’s Tau-b coefficient to measure the correlation between the optimal ordering and the ordering obtained using each method. Kendall’s Tau-b is a rank correlation coefficient adjusted for ties (Kendall, 1938) with values ranging between −1 and 1, with −1 signifying perfect inversion, 1 signifying perfect agreement, and 0 signifying the absence of association.

#### Default condition

ProACT dramatically increased the performance compared to random ordering according to all of our outcome measures (Fig. 2). Focusing on the transmissions per person measure, while the population mean was 0.05, the ProACT’s CMA was close to 0.15 for the top 1% of prioritized samples and gradually reduced to 0.1 for the top 10% (Fig. 2a). The top 1000 individuals in the ProACT ordering (3% of the population) transmitted 0.12 times (median across our 20 replicates), which was 2.4x higher than the median population average (Fig. 2c; see also Fig. S3 for numbers other than 1000). As desired, selecting fewer people from the top of ProACT prioritization resulted in more transmissions per person (Fig. 2a). Compared to optimal ordering, however, the adjusted score both increased and decreased as more individuals were selected (Fig. 2b). The adjusted metric shows that while ProACT substantially outperformed random ordering, it did not come close to the effectiveness that could be achieved using the (hypothetical) perfect ordering. The Kendall’s Tau-b correlation also showed a positive correlation between ProACT ordering and optimal ordering; although the correlation coefficient is far from perfect (Fig. 2d), the correlations are statistically significant in all replicates (*p* < 10^-9^; see Fig. S7a).

**Fig. 2.**
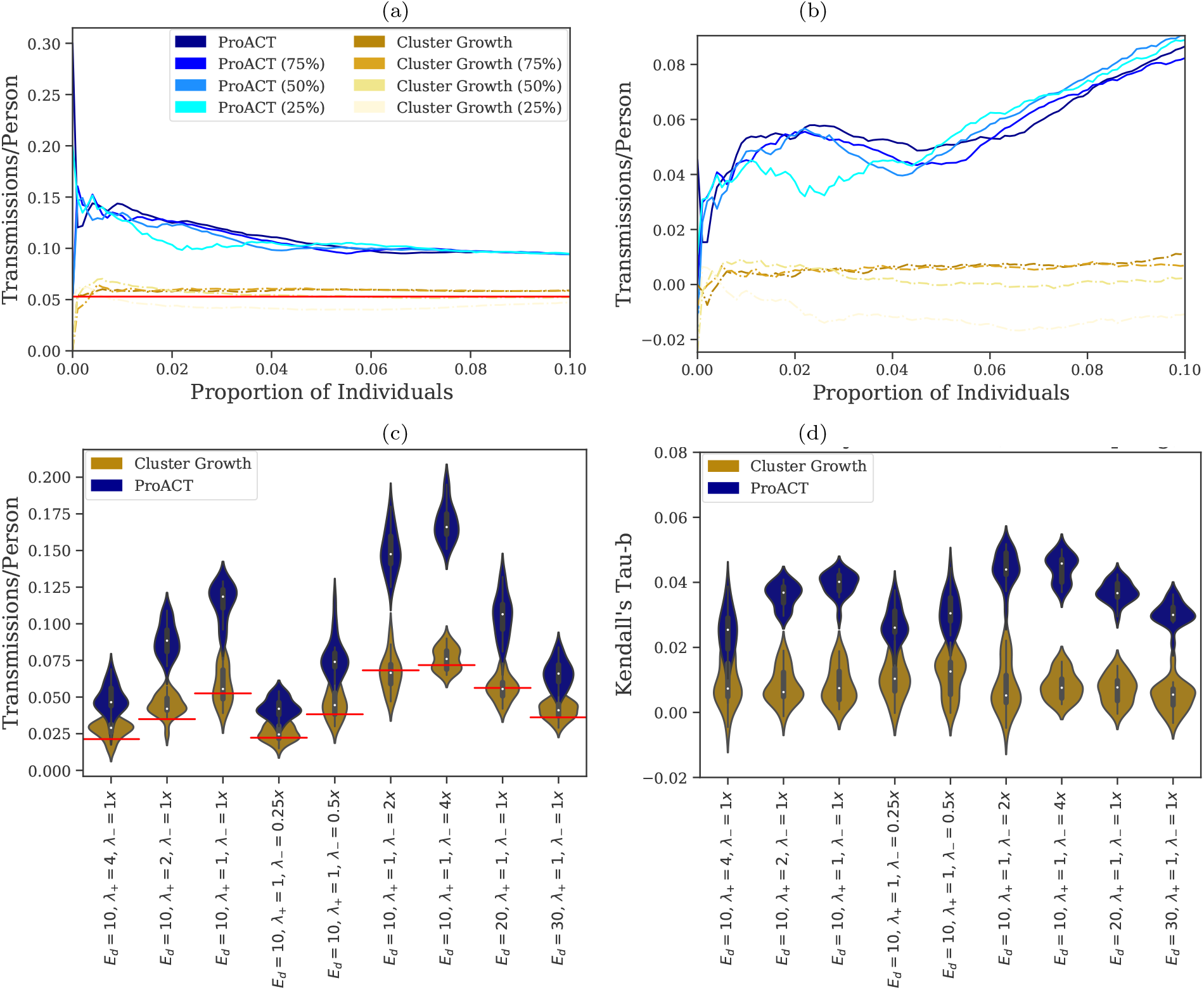
Effectiveness of prioritization on simulated datasets. The simulations were 10 years in length, prioritization was performed 9 years into the simulation, and the effectiveness of prioritization was computed during the last year of the simulation using four metrics (a-d). “Cluster Growth” denotes prioritization by inferring transmission clusters using HIV-TRACE at year 9 of the simulation and sorting clusters in descending order of growth rate since year 8. All curves were calculated using 20 simulation replicates. (a) Cumulative Moving Average (CMA) of the number of transmissions per person across the first decile of prioritized samples for the default simulation parameter set (see Fig. S4 for all model conditions, which show similar patterns.) The horizontal axis depicts the quantile of highest-prioritized samples (e.g. *x* = 0.01 denotes the top percentile), and the vertical axis depicts their average number of transmissions per person. Global average across all individuals (i.e., expectation under random ordering) is shown in red. The curves labeled with percentages denote subsampled datasets. (b) CMA of *adjusted* number of transmissions per person for the default model condition (See Fig. S5 for all model conditions, which show similar patterns.) For *adjusted* Transmissions/Person, 1 indicates the optimal ordering and 0 indicates random ordering. All other settings are similar to part a. (c) Average of the raw number of transmissions per person for the top 1000 individuals (see Fig. S3 for other counts) in a prioritized list vs. simulation parameter set (1000 individuals correspond to 1%–6% of all individuals across conditions). The violin plots are across 20 replicates and contain box plots with medians shown as white dots. Red horizontal lines show population mean (i.e., random prioritization). (d) Kendall Tau-b correlation between the optimal ordering of samples (i.e., based on their number of transmissions in year 10) and the orderings by the two prioritization methods. See Figure S6 for subsampled data. Distributions are across 20 replicates and are shown for each simulation condition.

Wertheim et al. (2018) have presented a method for prioritizing samples by clustering individuals based on viral genetic distance, tracking the size of each cluster over time, and prioritizing clusters in descending order of the growth rate. The approach can be extended to also order individuals (i.e., individuals belonging to clusters with high growth rates are prioritized higher; see Materials and Methods for details). ProACT consistently outperformed prioritization using cluster growth (Figs. 2). For example, the top 1000 individuals according to cluster growth transmitted on average to 0.06 other people, which, while higher than the population average, was half the 0.12 transmissions per person according to ProACT. Kendall-Tau results similarly indicate that ProACT has better correlation with the optimal ordering.

#### Impact of simulation parameters

We then tested the impact of three simulation parameters, namely the rate of stopping ART, the rate of starting ART, and the node degree in the sexual network (Figs. 2cd, S4, and S5).

As we increased the rate of stopping ART (λ_−_) (i.e., with lower adherence), the gap between ProACT and cluster growth grew. For example, the mean number of transmissions per person among the top 1,000 individuals chosen using ProACT and cluster growth were respectively 0.169 and 0.076 (a 1.21x improvement) for the condition with λ_−_ = 4x (Fig. 2c). This 1.21x improvement briefly increased to 1.26x and subsequently gradually decreased to 1.01x, 0.69x, and 0.63x as we reduced the rate or ART termination to 2x, 1x, 0.5x, and 0.25x. Kendall-Tau-b correlations show similar patterns (Fig. 2d); while almost all replicates of λ_−_ = 4x have *p* < 10^-20^, for the 0.25x case, all replicates have *p* > 10^-10^ and one of the replicates has *p* > 10^-3^ (Fig. S7a).

As we increased the rate of starting ART (λ_+_) (i.e., with faster diagnoses), as expected, the raw number of new infections caused per capita also reduced (Fig. 2c, S4a). While ProACT remained effective in finding high priority individuals, its performance compared to optimal ordering slightly degraded with higher λ_+_ (Figs. 2d and S5a). Also, the gap between ProACT and cluster growth decreased slightly. When observing the mean number of transmissions per person among the top 1,000 individuals chosen by each method (Fig. 2c), ProACT gave a 1.01x, 1.03x, and 0.71x improvement over cluster growth for λ_+_ set to 1x, 2x, and 4x, respectively.

Changing the expected number of sexual contacts per person (*E_d_*), which controls the speed of spread, did not have uniform effects (Figs. 2cd). Increasing *E_d_* from 10 to 20 did not substantially impact the performance of ProACT. However, for *E_d_* = 30, we observed a small but noticeable reduction in the performance of ProACT compared to the optimal ordering and cluster growth (Figs. 2d and S5d).

#### Impact of incomplete sampling

Subsampling the total dataset to include ¾, ½, or ¼ of all samples had only a marginal impact on the performance of ProACT according to the CMA metric (Figs. 2ab, S4, S5). Only at 25% sampling level did we observe a small reduction in the performance of ProACT compared to the optimal ordering. For example, with λ_+_ = 2x, ProACT’s performance remained quite similar across ≥ ½ sampling levels, but a reduction in performance was observed for the ¼ sampling level for both ProACT and cluster growth (Fig. S5a).

According to Kendall’s Tau-b, which measures the entire order not just the top individuals, there was a more noticeable degradation in performance due to sampling (Fig. S6). In particular, reduced sampling increased the *variance* across replicate simulations (note the wider distributions for reduced sampling in Fig. S6). Moreover, statistical significance of the correlations degrades with lower sampling (Fig. S7c-e). With ¼ sampling, unlike full sampling, many model conditions include *some* replicates where the ProACT ordering is not significantly better than random according to Kendall’s Tau-b.

#### Second order effects

We next asked if prioritization is effective in detecting people whose contacts also transmit abundantly. To do so, we explored a new outcome measure: the total number of transmissions from all contacts of a sample. Prioritizing samples whose contacts are likely to transmit can give public health officials a chance to find undiagnosed individuals (likely to transmit) through partner tracing from diagnosed individuals and to prioritize PrEP for uninfected individuals.

Across all model parameters, ProACT ordering outperformed random ordering and cluster growth according to the number transmissions per neighbor (Fig. 3). For example, contacts of the top 1000 individuals according to ProACT transmitted to 2.23 individuals on average (median across replicates), which is more than twice the number of transmissions by contacts across all individuals in the network (1.08). Just as with the previous outcome measure, advantages of ProACT over random prioritization or cluster growth were most pronounced for lower λ_+_ and higher λ_−_ (Fig. 3c). The Kendall Tau-b coefficients for the correlation between ProACT and the optimal ordering were high (Fig. S8); in fact, they were *higher* for the transmissions from contacts compared to transmissions from the prioritized person (e.g. median coefficient was 0.084 for contacts and 0.033 for the individuals in the default condition). These coefficients were highly significant across all models and sampling levels (Fig. S9a). Thus, ProACT was even more effective in finding individuals with active contact than it was for finding individuals who were not suppressed. These results were largely robust to reduced sampling, showing similar patterns of average performance but increased variance across replicates (Fig. S8 and S9c–e).

**Fig. 3.**
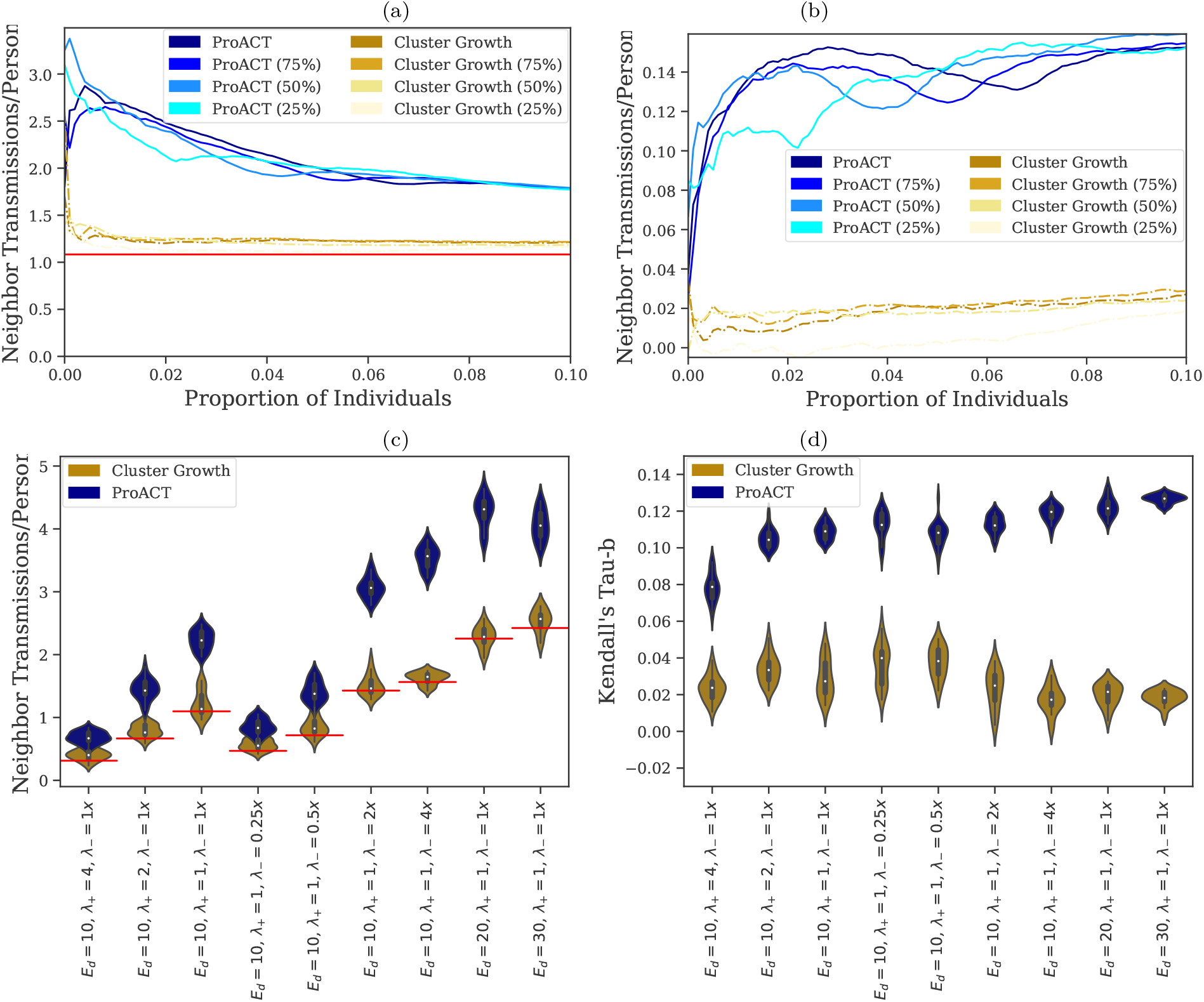
Second order effects. (a) CMA of the number of infections from contacts of the top individuals according to each ordering; other settings similar to Fig. 2a. (b) Similar to part (a) but adjusted for random and optimal ordering. (c) Number of transmissions from neighbours for the top 1000 individuals in a prioritized list vs. simulation parameter set. (d) Kendall Tau-b correlation between the number of contacts of each individual and their ordering by the two prioritization methods. See Figure S10 for subsampled data.

Further interrogating the properties of an individual and their ordering, we observed a substantial correlation between the number of contacts of samples in the sexual network and their position in the ProACT ordering (Fig. 3d). Thus, while ProACT only considers the phylogeny, it was able to prioritize those individuals that had high degrees in the sexual contact network (hidden to ProACT). These correlations were strongest for networks with high degree and weakest when the rate of diagnosis was very high. Reducing sampling did not substantially affect these results (Fig. S10).

### Real San Diego dataset

We next analyzed a dataset of 926 HIV-1 subtype B *pol* sequences obtained in San Diego between 1996 and 2018. To evaluate ProACT accuracy, we divided the data into deciles, with each decile defining two sets: *past* (sequences up to the decile) and *future* (sequences after the decile). We inferred a phylogeny from the sequences present in the *past* set using FastTree 2 Price et al. (2010), and we used ProACT to order all samples in this set. We then evaluated how the outcome measure correlates with the position of each individual in the ordering. We quantify the correlation using Kendall’s tau-b, a rank correlation coefficient adjusted for ties Kendall (1938). Values range between −1 and 1, with −1 signifying perfect inversion, 1 signifying perfect agreement, and 0 signifying the absence of association.

On real datasets, unlike the simulated data, the desired outcome measure, the number of new transmissions per person, is not known. Instead, we have to use inferred relationships. HIV-TRACE (used in our cluster growth approach) defines a pair of samples as “genetically linked” if their sequences are very similar (TN93 distance below 1.5%). We similarly use the TN93 sequence similarity as an outcome measure, but in addition to using a fixed threshold, we also use smoother functions (Fig. S11). We measure the number of linked individuals using a step function (1 if TN93 distance is below 1.5% and 0 otherwise) and an empirical smooth step function determined by fitting a mixture of three Gaussians to the distribution of pairwise TN93 distances (Material and Methods). We also explore an analytical smooth step function (parameterized sigmoid). Note that, when the step function is used, our outcome measure (computed for future transmissions) is exactly the same as what the cluster growth method uses for prioritizing (albeit, using past data). Thus, it is reasonable to expect the step function will favor cluster growth. As we move to smoother functions of distance to count genetic links, our measure is expected to become less biased in favor of HIV-TRACE.

Using both ProACT and cluster growth to prioritize individuals results in orderings of individuals with positive Kendall’s tau-b correlations to the number of future genetic links regardless of the time (i.e., decile) and the function used to count genetic links (Fig. 4). These correlations are statistically significant in almost all cases (Table 2 and Fig. 4). The correlation coefficient ranges ranges between 0.4 (ProACT; 10% time) and 0.1 (cluster growth; 20% time) for empirical function, and between 0.6 (cluster growth; 10% time) and 0.1 (ProACT; 80% time) for the step function.

**Fig. 4.**
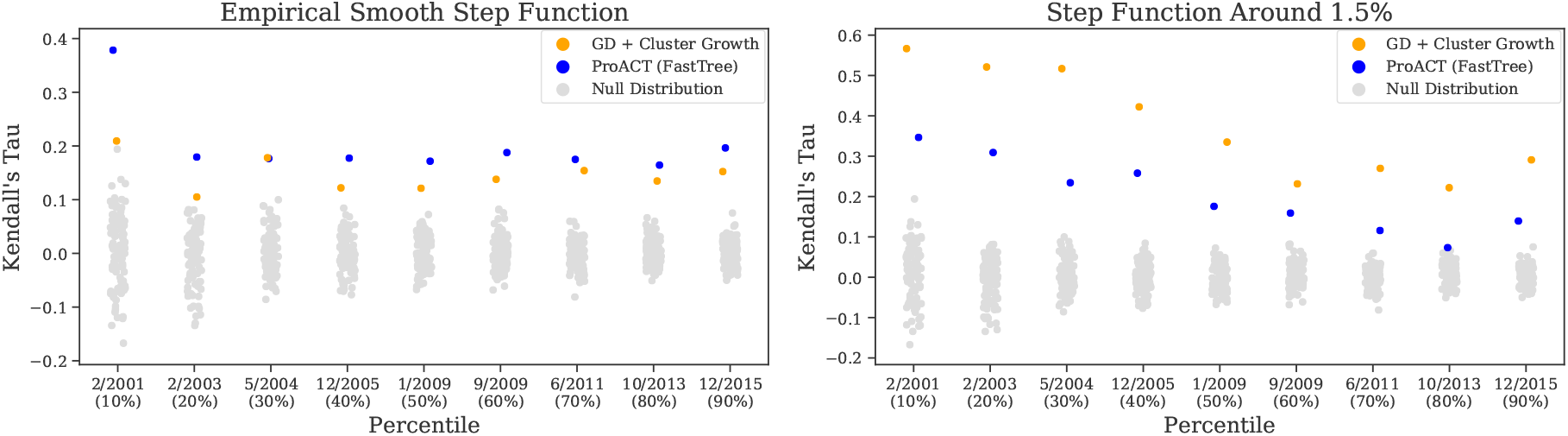
Kendall’s tau-b test results for ProACT ordering on real data using two score functions: an empirical smooth step function and a strict step function around 1.5%. The full San Diego dataset was split into two sets (*pre* and *post*) at each decile (shown on the horizontal axis). The individuals in pre were ordered using ProACT and by cluster growth, and they were given a “score” computed using a score function (see Materials and Methods). Kendall’s tau-b correlation coefficient was computed for each ordering with respect to the optimal possible ordering (i.e., sorting in descending order of the score). The null distribution was visualized by randomly shuffling the individuals in *pre*, and test *p*-values are shown in Table 2.

**Table 2.**
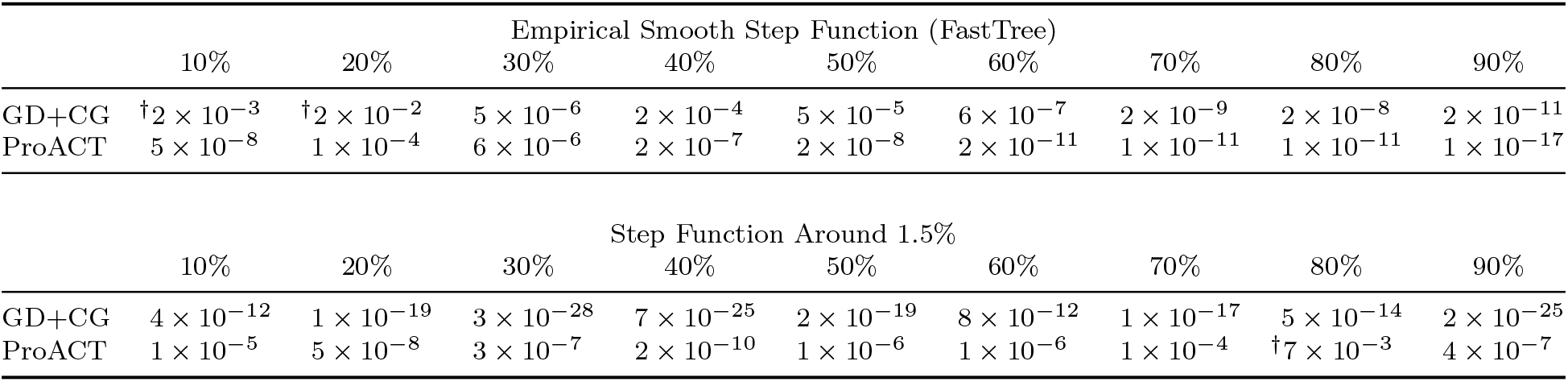
Kendall’s tau-b test for a null hypothesis that a given prioritization yields a total outcome measure no better than random. We show *p*-values for the real San Diego dataset for the first through ninth deciles using two outcome measure functions. Tests that failed to reject the null hypothesis with (uncorrected) *p*-value < 0.00138 (corresponding to *a* = 0.05 with a Bonferroni multiple hypothesis testing correct with *n* = 36) are marked with †.

|  |  | Empirical Smooth Step Function (FastTree) |  |  |  |  |  |  |  |
| --- | --- | --- | --- | --- | --- | --- | --- | --- | --- |
|  | 10% | 20% | 30% | 40% | 50% | 60% | 70% | 80% | 90% |
| GD+CG | † $2 \times 10^{-3}$ | † $2 \times 10^{-2}$ | $5 \times 10^{-6}$ | $2 \times 10^{-4}$ | $5 \times 10^{-5}$ | $6 \times 10^{-7}$ | $2 \times 10^{-9}$ | $2 \times 10^{-8}$ | $2 \times 10^{-11}$ |
| ProACT | $5 \times 10^{-8}$ | $1 \times 10^{-4}$ | $6 \times 10^{-6}$ | $2 \times 10^{-7}$ | $2 \times 10^{-8}$ | $2 \times 10^{-11}$ | $1 \times 10^{-11}$ | $1 \times 10^{-11}$ | $1 \times 10^{-17}$ |

|  |  | Step Function Around 1.5% |  |  |  |  |  |  |  |
| --- | --- | --- | --- | --- | --- | --- | --- | --- | --- |
|  | 10% | 20% | 30% | 40% | 50% | 60% | 70% | 80% | 90% |
| GD+CG | $4 \times 10^{-12}$ | $1 \times 10^{-19}$ | $3 \times 10^{-28}$ | $7 \times 10^{-25}$ | $2 \times 10^{-19}$ | $8 \times 10^{-12}$ | $1 \times 10^{-17}$ | $5 \times 10^{-14}$ | $2 \times 10^{-25}$ |
| ProACT | $1 \times 10^{-5}$ | $5 \times 10^{-8}$ | $3 \times 10^{-7}$ | $2 \times 10^{-10}$ | $1 \times 10^{-6}$ | $1 \times 10^{-6}$ | $1 \times 10^{-4}$ | † $7 \times 10^{-3}$ | $4 \times 10^{-7}$ |

The comparison between ProACT and cluster growth depends on the choice of the function to count links. When counting the number of links using the step function, prioritization by cluster growth consistently outperforms ProACT for all deciles of the dataset. These results are not surprising, given that we count HIV-TRACE links both to prioritize and to evaluate. However, according to the empirical smooth step function learned from the TN93 distances, ProACT outperforms cluster growth in all except one time point, where they are tied.

To further test whether the smoothness of the link-counting function applied to TN93 distances is a factor in deciding the relative accuracy of methods, we used a sigmoid function to replace the step function while keeping the inflection point at 1.5% (Fig. S11). We observed that as the outcome measure function becomes more smooth, ProACT’s performance improves with respect to prioritization by cluster growth (Fig. 5, Table S1). Based on the more smooth sigmoid function (λ = 5), ProACT outperforms cluster growth in all but one case where they are tied. Thus, simply counting distances close to 1.5% as partial links leads to evaluations that favor ProACT.

**Fig. 5.**
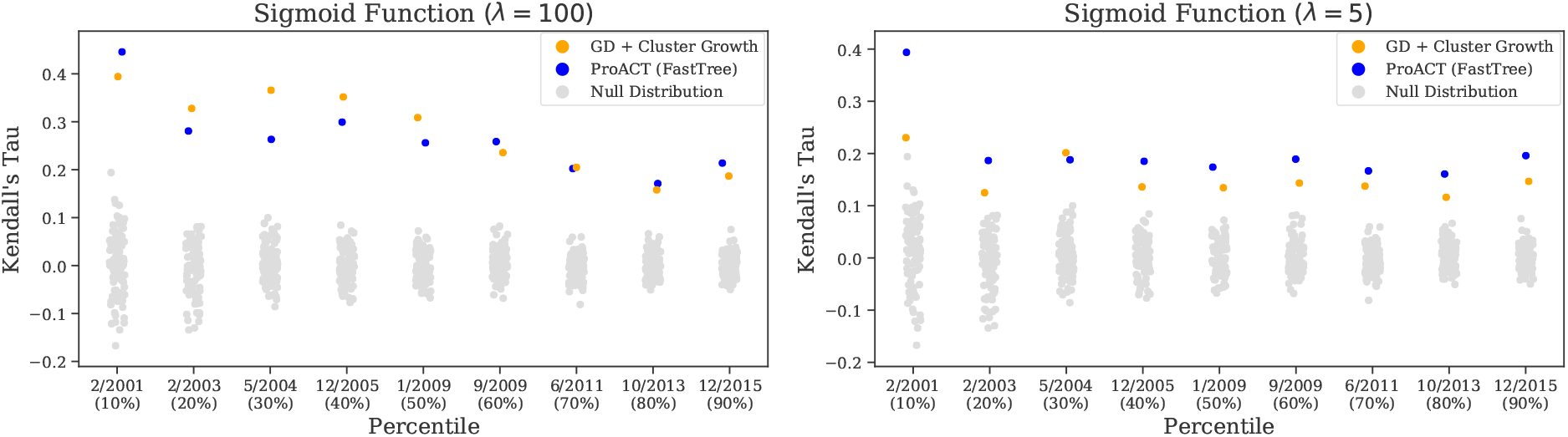
Kendall’s tau-b test results for ProACT ordering on real data using the sigmoid score functions with λ = 100 and λ = 5. The full San Diego dataset was split into two sets (*pre* and *post*) at each decile (shown on the horizontal axis). The individuals in pre were ordered using ProACT and by cluster growth, and they were given a “score” computed using a score function (see Materials and Methods). Kendall’s tau-b correlation coefficient was computed for each ordering with respect to the optimal possible ordering (i.e., sorting in descending order of the score). The null distribution was visualized by randomly shuffling the individuals in *pre*, and test *p*-values are shown in Table S1.

As time increases, both methods experience seemingly downward trends in their tau coefficients, but the null distribution of tau coefficients also tightens (Fig. 4). Thus, both methods consistently do significantly better than expected by random chance and there is no clear relationship between *p*-values of individual tool and time (Table 2). However, both for the step function and the sigmoid functions, ProACT’s relative performance with respect to cluster growth tends to improved over time.

## Discussion

We start by discussing observed results and then comment on practical implications of this paper both for public health and for future research in molecular epidemics.

### Discussion of Results

In our simulations, ProACT was least effective in conditions with very low rate of ART termination, which correspond to very high adherence, or high rates of ART initiation. As expected, the total number of new infections originated from samples is low when adherence is high (Fig. S4) reducing the opportunity for improving the ordering. Thus, ProACT is most beneficial in settings where termination of ART or late diagnosis lead to individuals who transmit frequently.

ProACT was quite robust to impacts of subsampling individuals and only at ¼ sampling did we start to lose accuracy. We remind the reader that a ¼ sampling does not mean that ¼ of all infected individuals are in the dataset. Rather, it means that ¼ of diagnosed individuals are available to us. Recall that, in our model, diagnosed individuals are immediately sequenced and put on ART (which they may or may not sustain). At any point in time, a large partition of individuals who are infected are not diagnosed and thus not sampled. In other words, the full sampling case should not be misunderstood as including undiagnosed people. Rather, lack of full sampling corresponds to a case where some samples are known to *some* clinic but are not included in the study, perhaps due to a lack of sequencing or data sharing.

ProACT far outperformed random ordering. However, we note that, despite the strong performance, there is much room left for future improvement: ProACT consistently ranges in its outcome measure between 2% to 8% of the theoretical optimal value when selecting up to 10% of top-priority samples. Thus, there is great room for improvement in identifying high-value individuals. It will be unrealistic to expect that any statistical method based solely on sequence data (and perhaps also commonly available metadata, e.g. sampling times) will be able to come close to the optimal ordering. Nevertheless, it remains likely that methods better than ProACT could in fact be developed. Moreover, here, we used ML methods to infer trees and used mutation rate branch lengths. We made these choices mostly for computational expediency. However, ProACT algorithm can be applied on the potentially more accurate Bayesian estimates of the phylogeny. Also, one can attempt to use ProACT after dating the tree. Whether either adjustment results in substantial improvements should be studied in the future.

### Implications of Results

We formalized a useful approach for thinking about the effectiveness of public health intervention in molecular epidemics. Instead of focusing on the accuracy of methods of reconstructing phylogenetic trees or transmission networks, a question fraught with difficulties, we asked a more practical question. Given molecular epidemic data, can the methods, whether phylogenetic or clustering-based, prioritize samples for increased attention by public health? Using molecular epidemics for prioritization is, of course, not a new idea. For example, Wertheim et al. (2018) presented a method to prioritize samples based on the growth rates of their transmission clusters. Vasylyeva et al. (2018) performed a phylogeographic analysis to reconstruct HIV movement among different locations in Ukraine in order to infer region-level risk prioritization. Much earlier even, Mellors et al. (1996) predicted HIV patient prognosis by quantifying HIV RNA in plasma; predicted prognosis can subsequently be used as a prioritization rank. However, we hope that our formal definition of the problem as a computational question (i.e., prioritization), in addition to our extensive simulations and developed metrics of evaluation, will stir further work in this area. As stated before, it seems likely that more advanced methods than our simple prioritization approach can improve performance beyond ProACT in the future.

ProACT prioritizes individuals, not clusters. Prioritizing treatment followup or partner tracing for individuals based on their perceived risk of future transmission promises to be perhaps more effective than targeting clusters. However, such targeted approaches also pose ethical questions that have to be considered. For example, we may not want the algorithm to be biased towards particular demographic attributes. ProACT does not use *any* metadata in its prioritization, reducing risks of such biases. It simply uses the viral phylogeny. Nevertheless, it is possible that factors such as the depth of the sampling of a demographic group can in fact change branch length patterns in the phylogeny and make ProACT less or more effective for certain demographic groups. These broader implications of individual prioritization and impacts of demographics on the performance of ProACT should be studied more carefully in future.

The main practical question is what can be done with a prioritized list of known samples. We mentioned that using followups, public health officials can try to ensure sustenance of ART for prioritized individuals, and using partner tracing, they can target PrEP and HIV testing to contacts of prioritized individuals. Followups, PrEP, and targeted testing are all expensive and can benefit from prioritization. Interestingly, our results indicated that ProACT ordering is a function of features of the sexual contact. For example, we showed that ProACT orders correlate with the degree of nodes in the sexual network. These results are significant given the fact that ProACT is given no direct data the sexual network. The fact that ProACT captures (contact) network features means that even if a prioritized sample is already on ART (and thus unlikely to transmit), his/her sexual contacts can be good targets for interventive care.

One may wonder whether ordering by branch lengths will result in orderings that fail to change with time and reflect the changes in the epidemic. To answer this question, on the San Diego PIRC data, we asked how fast the ProACT ordering changes as time progresses. To do so, we computed Kendall’s tau-b correlations to the ProACT ordering obtained using only the first decile of the dataset (Fig. S12). There was a strong but diminishing correlation with the initial ordering. The correlations started at 1 (as expected) and gradually decreased in the ninth decile to 0.522. The results show that as desired, ProACT orders do in fact change with time, albeit gradually. The gradual change implies that certain individuals remain high-priority as time progresses. In practical use, ProACT ordering should be combined with clinical knowledge about the status of individual patients. For example, high priority individuals according to ProACT can be given lower priority if they manage to constantly remain suppressed with multiple followups. More broadly, the ProACT ordering should be considered one more tool for prioritizing clinical care, but valuable clinical knowledge, not incorporated into the algorithm, should also be exploited.

Finally, a question faced by public health officials is whether the cost of targeting diagnosed individuals for followups and partner tracing is worth the reduction in future cases. The answer to that question will inevitably depend on who is targeted. For example, in our default simulation case, targeting individuals randomly can directly prevent 0.053 transmissions per chosen person in the next 12 months, whereas targeting top 1000 individuals according to ProACT would directly target 0.115 transmissions. Thus, prioritization can in fact change the cost-benefit analyses. Moreover, given a prioritization, one can use simulations to predict the outcome measure for the top individuals (similar to Fig. S5) and use metrics such as quality-adjusted life-year (QALY) to estimate how many top individuals should be targeted for the cost to justify the benefits.

## Materials and Methods

### Simulated Datasets

We use FAVITES to simulate a sexual contact network, transmission network, viral phylogeny, and viral sequences emulating HIV transmission in San Diego from 2005 to 2014 (Moshiri et al., 2018).

Transmissions are modeled using a compartmental epidemiological model with 5 states: Susceptible (S), Acute HIV Untreated (AU), Acute HIV Treated (AT), Chronic HIV Untreated (CU), and Chronic HIV Treated (CT). Individuals in state S (i.e., uninfected) can only transition to state AU. Each infected state *x* ∈ {AU, AT, CU, CT} defines a “rate of infectiousness” λ_S,*x*_: given an uninfected individual *u* in state S who has *n_x_* sexual partners in state *x* ∈ {AU, AT, CU, CT}, the transition of *u* from S to AU is a Poisson process with rate λ_*u*_ = Σ_*x*∈{AU,AT,CU,CT}_ *n_x_*λ_S,*X*_. To mimic reality, where ART significantly reduces the risk of transmission, rates are chosen such that λ_S,AU_ > λ_S, CU_ > λ_S, AT_ > λ_S, CT_ ≈ 0. At the beginning of the epidemic simulation, all initially uninfected individuals are placed in state S, and all initially infected (i.e., “seed”) individuals are distributed among the 4 infected states according to their steady-state proportions. This model is a simplified version of the model proposed by Granich et al. (2009).

Once the transmissions and sample times are obtained, the viral phylogeny evolves inside the transmission tree under a coalescent model of evolution with logistic within-host viral population growth and a bottleneck event at the time of transmission (i.e., initial viral population size is 1) (Ratmann et al., 2017). This process produces a separate viral phylogeny for each seed individual, so we also need a tree for seed individuals. Each *seed* individual of the epidemic is the root of an independent viral phylogeny, and these trees were merged by simulating a seed tree with one leaf per seed node under a non-homogeneous Yule model (Le Gat, 2016) with rate function λ(*t*) = *e*^−*t*^2^^+ 1 scaled to have a height of 25 years to match the estimate of the time of the most recent common ancestor of HIV in San Diego (Moshiri et al., 2018). A mutation rate was sampled for each branch independently from a truncated normal random variable from 0 to infinity with a location parameter of 0.0008 and a scale parameter of 0.0005 to scale branch lengths from years to expected number of per-site mutations (Moshiri et al., 2018).

For the most part, we use the base parameters used in Moshiri et al. (2018) that sought to model the San Diego HIV epidemic from 2005 to 2014, with the following modifications to better capture reality. See Table S2 for the full set of parameters of the default condition.

#### Sexual contact network

To capture the scale-free nature of the sexual contact network, Moshiri et al. (2018) used the Barabàsi-Albert (BA) model (Barabási and Albert, 1999). In addition to the scale-free property, in HIV sexual networks, we typically observe many densely-connected communities Rothenberg et al. (1998), a property the BA model fails to directly model. To have control over the number of communities, we simulated sexual contact networks such that networks contained 20 BA communities, each with 5,000 individuals. In the base condition, the expected degree of connection between an individual and somebody *within* their community was chosen to be 10, and the expected degree between an individual and somebody *outside* their community was chosen to be 1. Each community was simulated separately using the BA model and connections between communities were chosen uniformly at random, akin to the Erdős-Rényi model (Erdos and Rényi, 1959). Estimates from the literature put the number of contacts at 3–4 during a single year (Rosenberg et al., 2011). Because our simulated sexual contacts remain static over the 10 year simulation period, we explore mean degrees between 10 and 30.

#### Epidemic initialization

In Moshiri et al. (2018), at the start of the epidemic, all infected individuals were in state AU. Here, instead, we randomly distribute initially infected individuals according to expected proportions of the states. To find these proportions, we ran simulations in which all seed individuals were in state AU, and we observed the proportion of individuals in each state over time, which reached a steady-state fairly early in the simulations (Fig. S13).

#### Time of sequencing

In Moshiri et al. (2018), viral sequences are obtained from individuals exactly at the end time of the 10-year simulation period. In reality, however, HIV patients are typically sequenced when they first visit a clinic to receive ART. Thus, it is expected that the terminal branch lengths of trees simulated in Moshiri et al. (2018) are artificially longer than would be expected. Instead, we sample viral sequences from individuals the first time they begin ART (i.e., the first time they enter state AT or CT). Our current simulation better captures standards of care in advanced health care systems.

#### Simulated data analysis

For each simulated sequence dataset, using FastTree 2 (Price et al., 2010), a phylogenetic tree was inferred under the GTR+Γ model from the sequences obtained in the first 9 years of the simulation. These trees were then MinVar-rooted using FastRoot (Mai et al., 2017), and ProACT was run on the resulting trees.

### PIRC San Diego Dataset

To test ProACT on real data, we used a Multiple Sequence Alignment (MSA) of 926 HIV-1 subtype B *pol* sequences from San Diego collected by the UC San Diego Primary Infection Resource Consortium (PIRC). PIRC is one of the largest longitudinal cohorts of samples in the United States. By design, PIRC strives to include acute infections (as much as 40% of recruited individuals are during acute or early stages of infection). Access to the data was obtained through a proposal submitted to PIRC.

A phylogenetic tree was inferred from the MSA under the GTR+Γ model using FastTree 2 (Price et al., 2010), and the resulting tree was MinVar-rooted using FastRoot (Mai et al., 2017). For each decile, using TreeSwift (Moshiri, 2018), the full tree was pruned to only contain samples obtained up to the end of that decile. ProACT was run on each of the resulting trees.

### Evaluation Procedure

#### Simulated data

To measure the efficacy of a given ProACT selection, because the true transmission histories are known in simulation, we simply average the number of infections caused by the individuals in the selection in the last year of simulation (i.e, after prioritization) to obtain a raw outcome measure.

Let *A* = {1,…, *n*} denote the first,…, *n*-th sampled individual in the current time step (years 1-9 in our simulations). For each individual *i*, let *c*(*i*) denote the number of individuals directly infected by *i* in the next time step (year 10 in our simulations). Given any set of individuals *s* ⊆ *A*, let 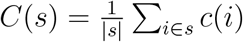 denote the average *c*(*i*) for all individuals *i* ∈ *s*.

Let *x* = (*x*_1_,…,*x_n_*) denote an ordering of *A*. The (unadjusted) Cumulative Moving Average (CMA) of *x* up to *i* is *C* ({*x*_1_,…, *x_i_*}). Let *o* = (*o*_1_,…, *o_n_*) denote the ordering of *A* in which elements are sorted in descending order of *c*(*i*) (i.e., the optimal ordering), with ties broken arbitrarily. We defined the adjusted CMA of *x* up to *i* as

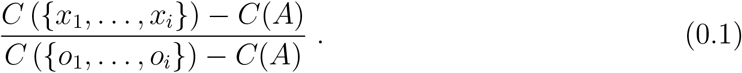

We use Equation 0.1 to measure the effectiveness of a selection of the top *i* individuals from each ordering of all individuals. We explore *i* for 1 to 10% of the total number of samples (i.e., 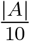).

#### Real data

The sequences were sorted in ascending order of sample time and, for each decile, they were split at the decile to form two sets: *pre* and *post*. A phylogenetic tree was inferred from the sequences in *pre* under the GTR+Γ model using FastTree 2 (Price et al., 2010) and MinVar-rooted (Mai et al., 2017). Using the resulting tree, ProACT ordered the samples. Then, pairwise distances were computed between each sequence in *pre* and each sequence in *post* under the Tamura-Nei 93 (TN93) model (Tamura and Nei, 1993) using the `tn93` tool of HIV-TRACE (Kosakovsky Pond et al., 2018).

A natural function to compute the risk of a given individual *u* in *pre*, similar to that proposed by Wertheim et al. (2018), is to simply count the number of individuals in *post* who are genetic links to *u*, i.e., Σ_*v*∈*post*_ [*d*(*u,v*) ≤ 1.5%]. In other words, the score function is simply a step function with value 1 for all distances less than or equal to 1.5% and 0 for all other distances. However, the selection of 1.5% as the distance threshold, despite being common practice in many HIV transmission clustering analyses, is somewhat arbitrary, and a step function exactly at this threshold may be overly strict (e.g. should a pairwise distance of 1.51% be ignored?).

To generalize this notion of scoring links, we utilized three analytical score functions. The first is the aforementioned step function *f*_1_(*d*) = [*d* ≤ 1.5%]. The second is a sigmoid function 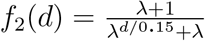 with the choice of λ = 100 and λ = 5 (Fig. S11). The third is an empirical scoring function learnt from the data by fitting a mixture model of three Gaussian random variables onto the distribution of pairwise TN93 distances 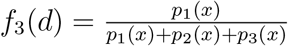, where *p*_1_ (*x*) is the Probability Density Function (PDF) of the Gaussian component with smallest mean and *p*_2_ (*x*) and *p*_3_ (*x*) are the remaining Gaussian components (Fig. S11). Specifically, the three Gaussian fits were parameterized by (*μ*_1_=0.0191, *σ*_1_=0.0103), (*μ*_2_=0.0609, *σ*_2_=0.0118), and (*μ*_3_=0.118, *σ*_3_=0.0468), respectively.

For each of these function, for each decile to define *pre* and *post*, we performed a Kendall’s tau-b test to compare the prioritization approaches (Kendall, 1938). To generate a null distribution in Figure 4, we randomly shuffled the individuals in *pre* repeatedly; note however that the *p*-values reported in Table 2 are the theoretical *p*-values computed by the tau-b test, not empirically estimated from our repeated shuffling.

## Acknowledgements

We thank Susan B. Little for providing the San Diego HIV sequence dataset used in this study. We also thank Joel O. Wertheim and Sanjay R. Mehta for fruitful discussions that helped motivate the development of ProACT.

This work was supported by the National Institutes of Health (5P30AI027767-28, AI100665, AI106039, and MH100974) and a developmental grant from the University of California, San Diego Center for AIDS Research (P30 AI036214), supported by the National Institutes of Health.

