## Supplementary Materials for "HIV Care Prioritization using Phylogenetic Branch Length"

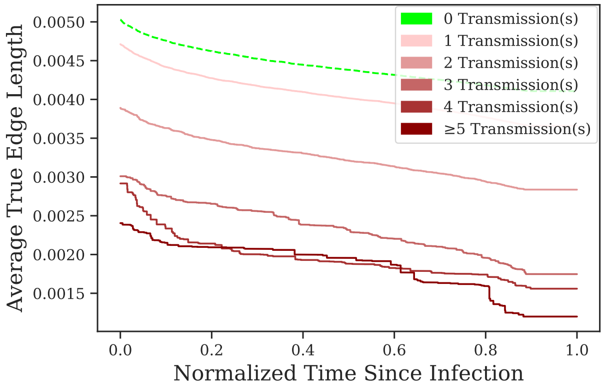


Fig. S1. As time progresses, the true incident branch length of each individual tends to decrease. This holds in inferred phylogenies as well (Fig. 1d).

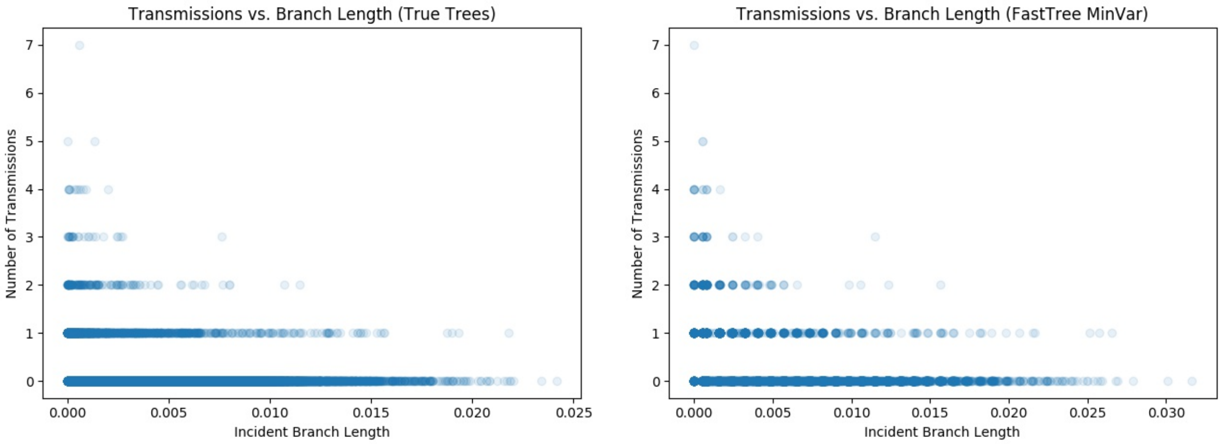


Fig. S2. Number of transmissions vs. incident branch lengths for individuals in a simulated epidemic. The epidemic was run for 10 years, samples were obtained at the 9-year mark, and a phylogeny was inferred using FastTree 2 Price et al. (2010) and subsequently MinVar-rooted Mai et al. (2017). Number of transmissions were measured between the 9-year and 10-year mark.

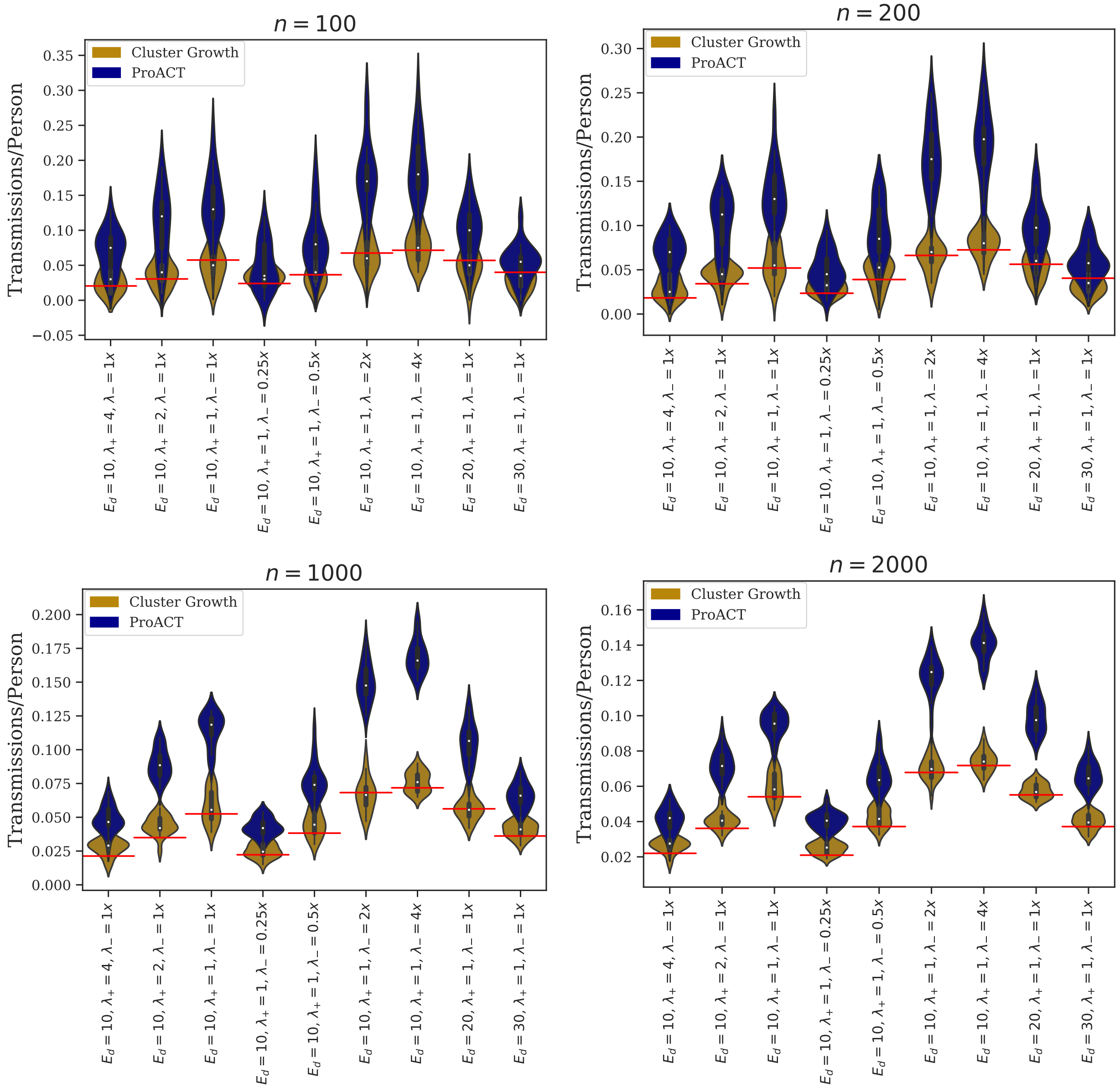

Fig. S3. Efficacy on datasets simulated using FAVITES. Average of the raw number of transmissions per person for the top $n$ individuals in a prioritized list vs. simulation parameter set across various values of $n$. The violin plots depicted are across 20 replicates and contain box plots with distribution medians shown as white dots and distribution means shown as dashed grey lines.

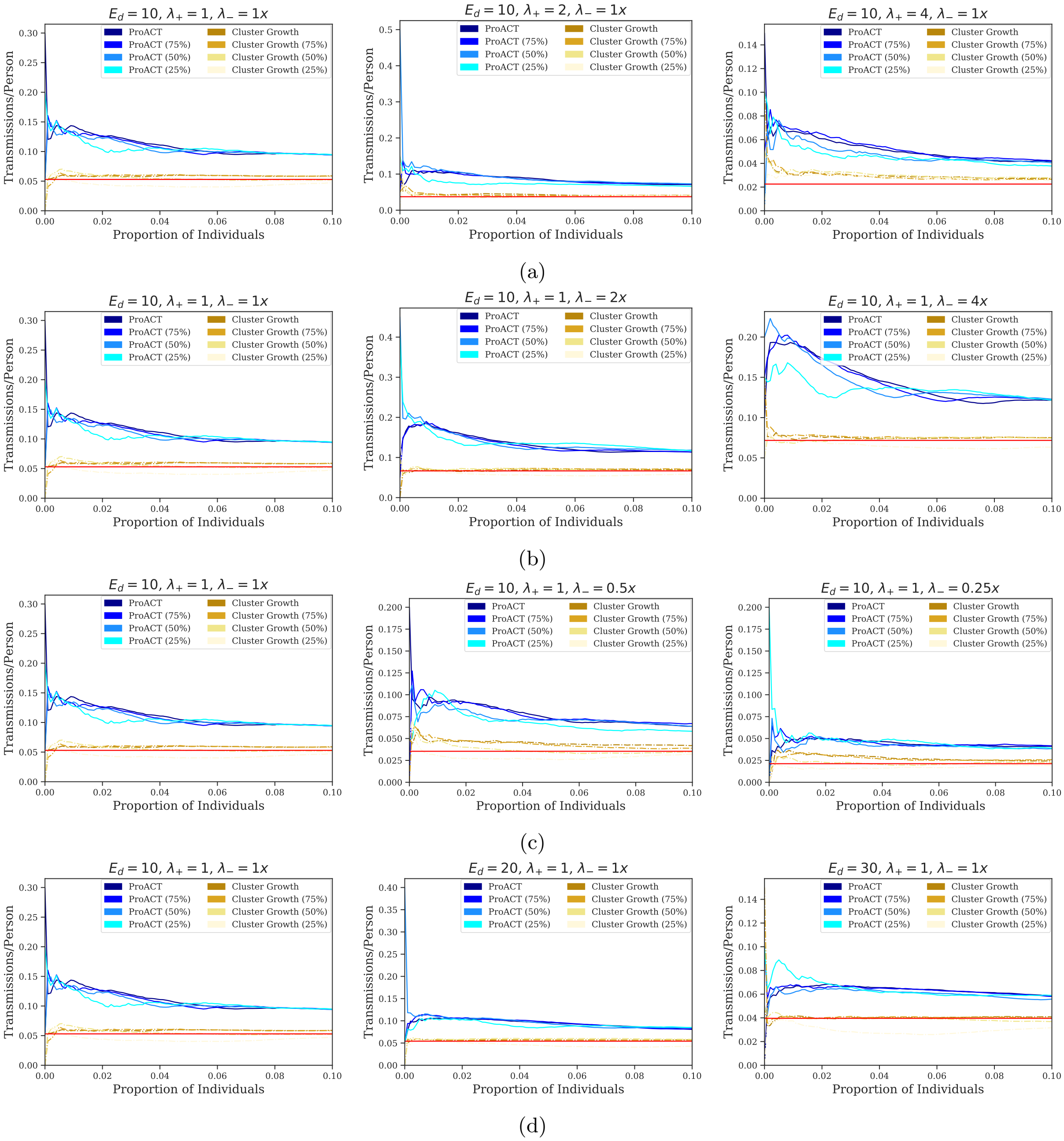


Fig. S4. Efficacy on datasets simulated using FAVITES. Cumulative Moving Average (CMA) of number of transmissions per person across all sample for each simulation parameter set.

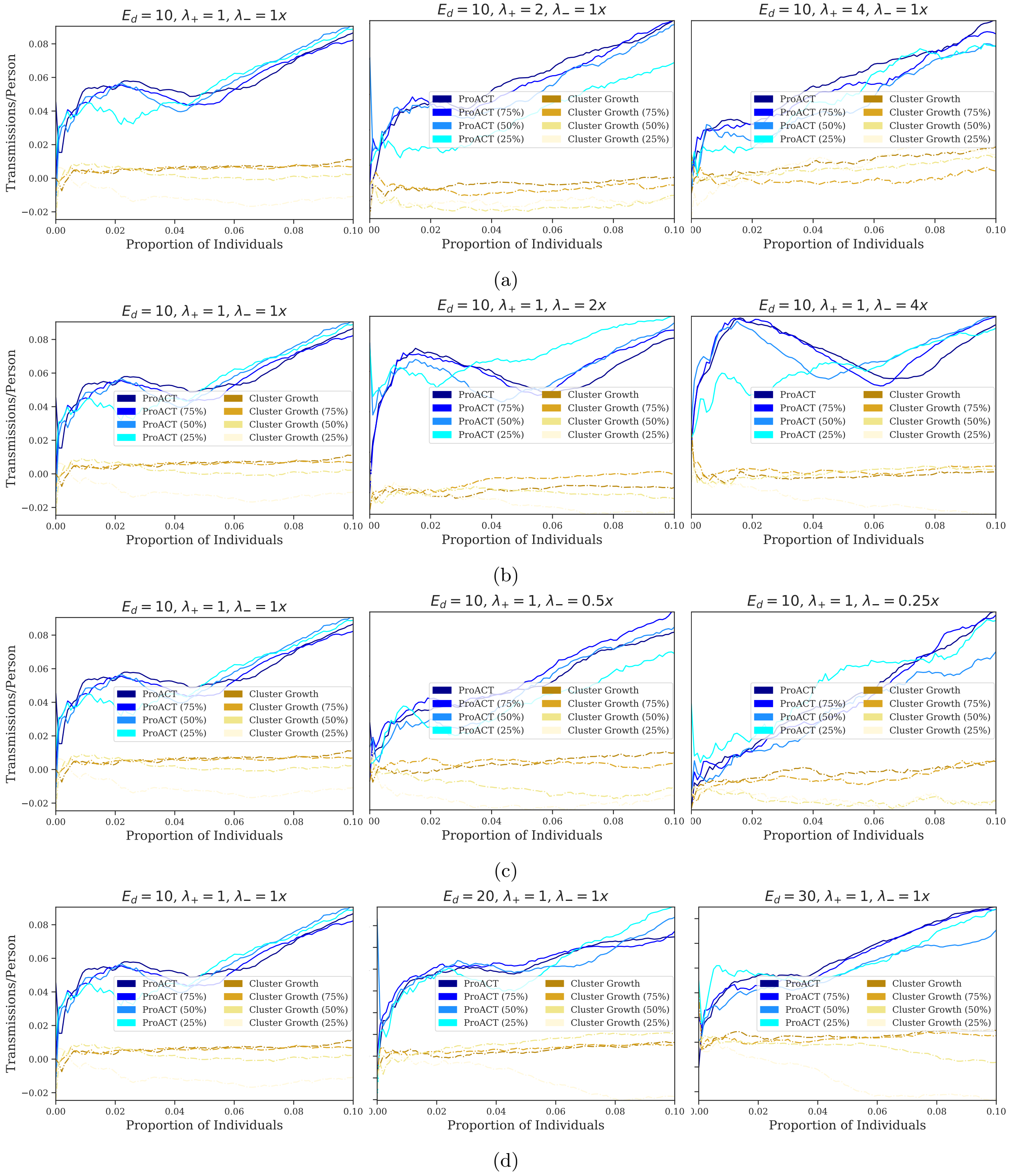


Fig. S5. ProACT performance on datasets simulated using FAVITES. Cumulative Moving Average (CMA) of adjusted number of transmissions per person across the first decile of prioritized sample for each simulation parameter set. The horizontal axis depicts the quantile of highest-prioritized sample (e.g. $x = 0.01$ denotes the top percentile), and the vertical axis depicts their adjusted average number of transmissions per person (1 indicates the optimal ordering, and 0 indicates an ordering that is no better than random). In our simulations, we varied three parameters of interest: (a) the rate of ART initiation ($\lambda_+$), (b-c) the rate of ART termination ($\lambda_-$), and (d) the expected degree of the sexual network ($E_d$). The simulations were 10 years in length, prioritization was performed 9 years into the simulation, and the adjusted average number of transmissions per person was computed during the last year of the simulation. The curves labeled "Cluster Growth" denote prioritization by inferring transmission clusters using HIV-TRACE at year 9 of the simulation and sorting clusters in descending order of growth rate since year 8. The curves labeled with percentages denote subsampled datasets. All curves were calculated using 20

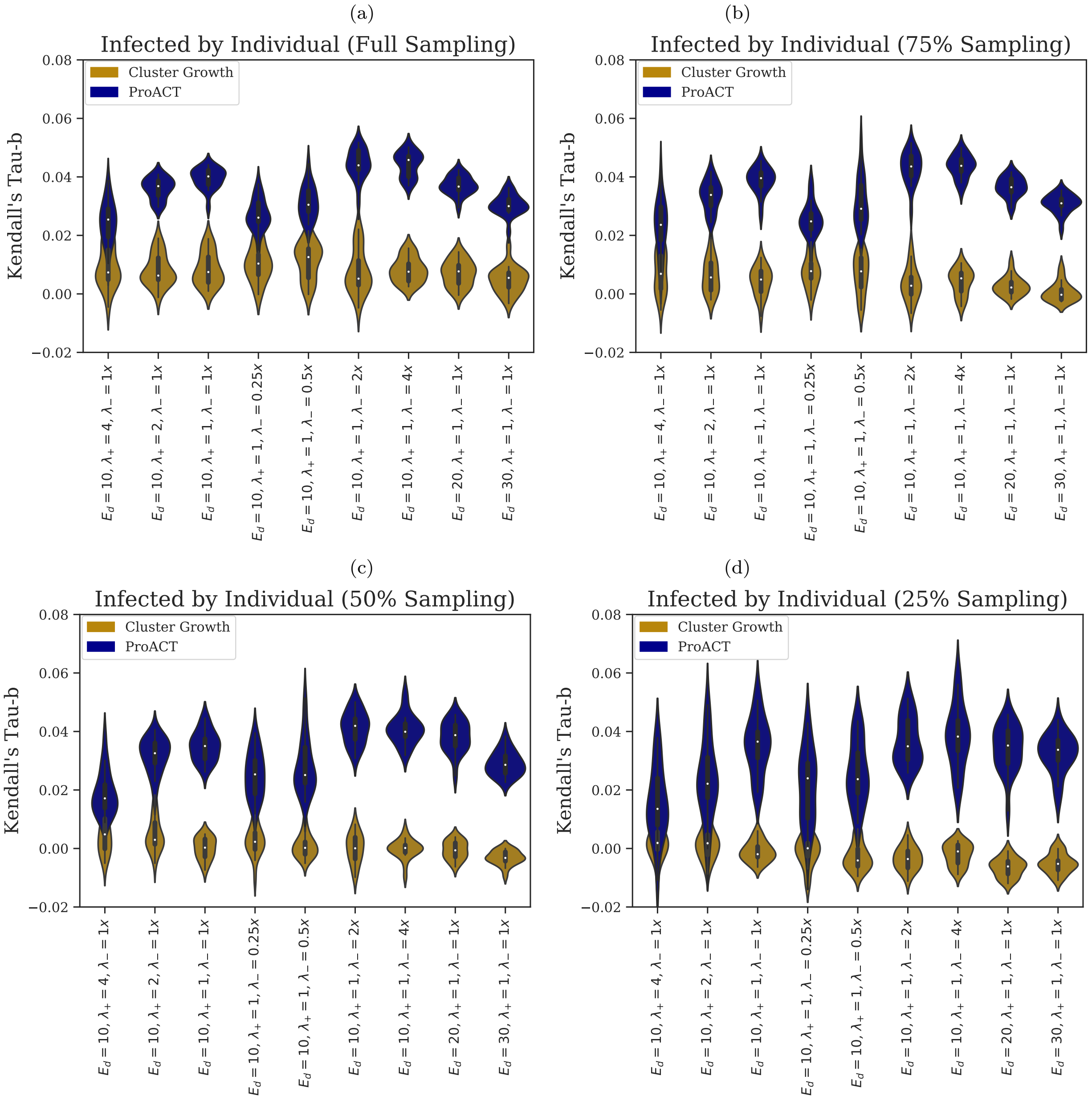


Fig. S6. Kendall Tau-b correlation between the optimal ordering of samples (i.e., based on their number of transmissions in year 10) and the orderings by the two prioritization methods, where effectiveness is measured by counting the number of future transmissions by a given individual. Distributions are across 20 replicates and are shown for each simulation condition. Results are shown for full sampling and subsampled data.

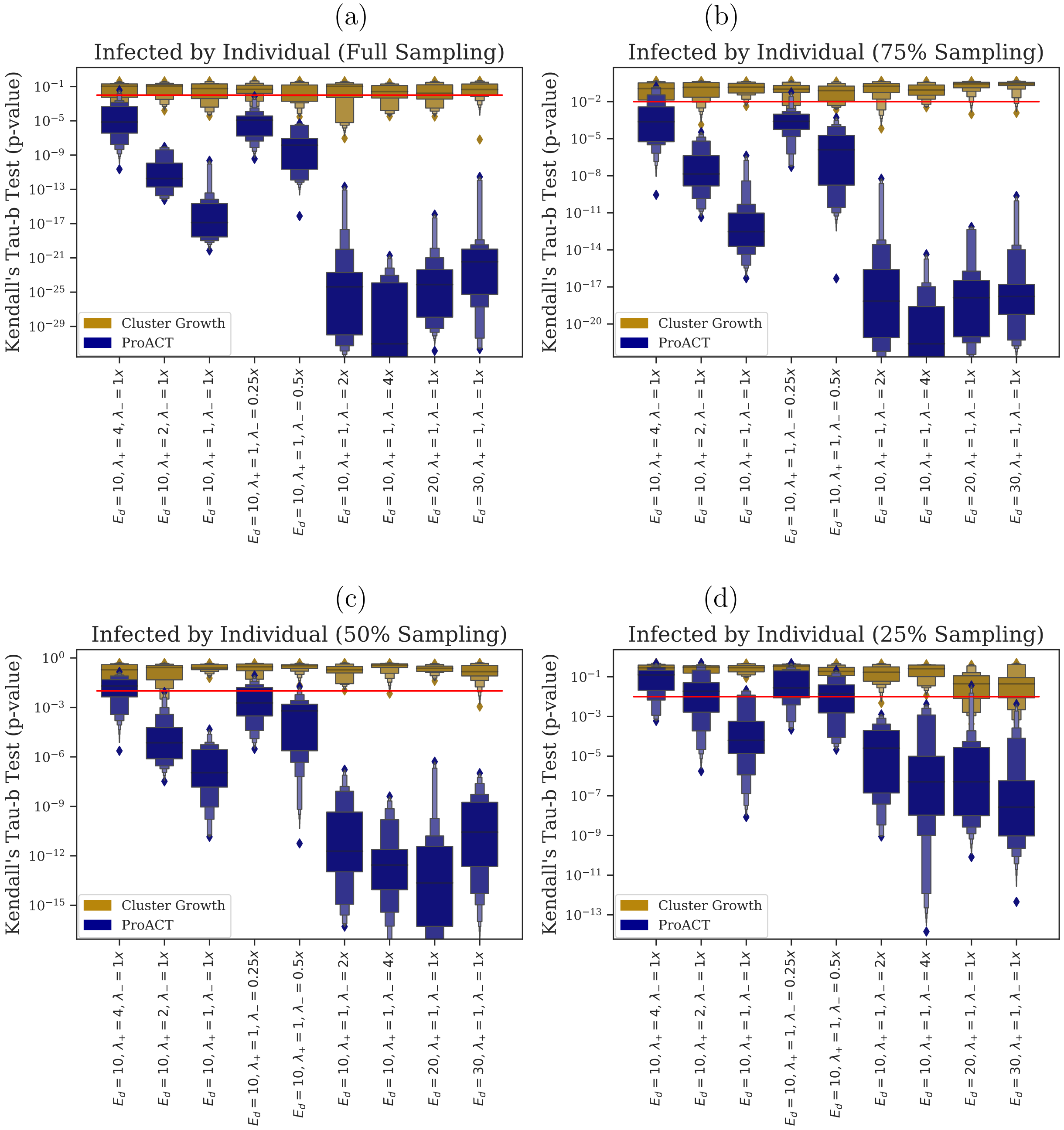


Fig. S7. $p$-values of one-sided Kendall Tau-b correlation tests between the optimal ordering of samples (i.e., based on their number of transmissions in year 10) and the orderings by the two prioritization methods, where effectiveness is measured by counting the number of future transmissions by a given individual. Distributions are across 20 replicates and are shown for each simulation condition. Results are shown for full sampling and subsampled data. See Figure S6 for Tau-b test statistic distributions.

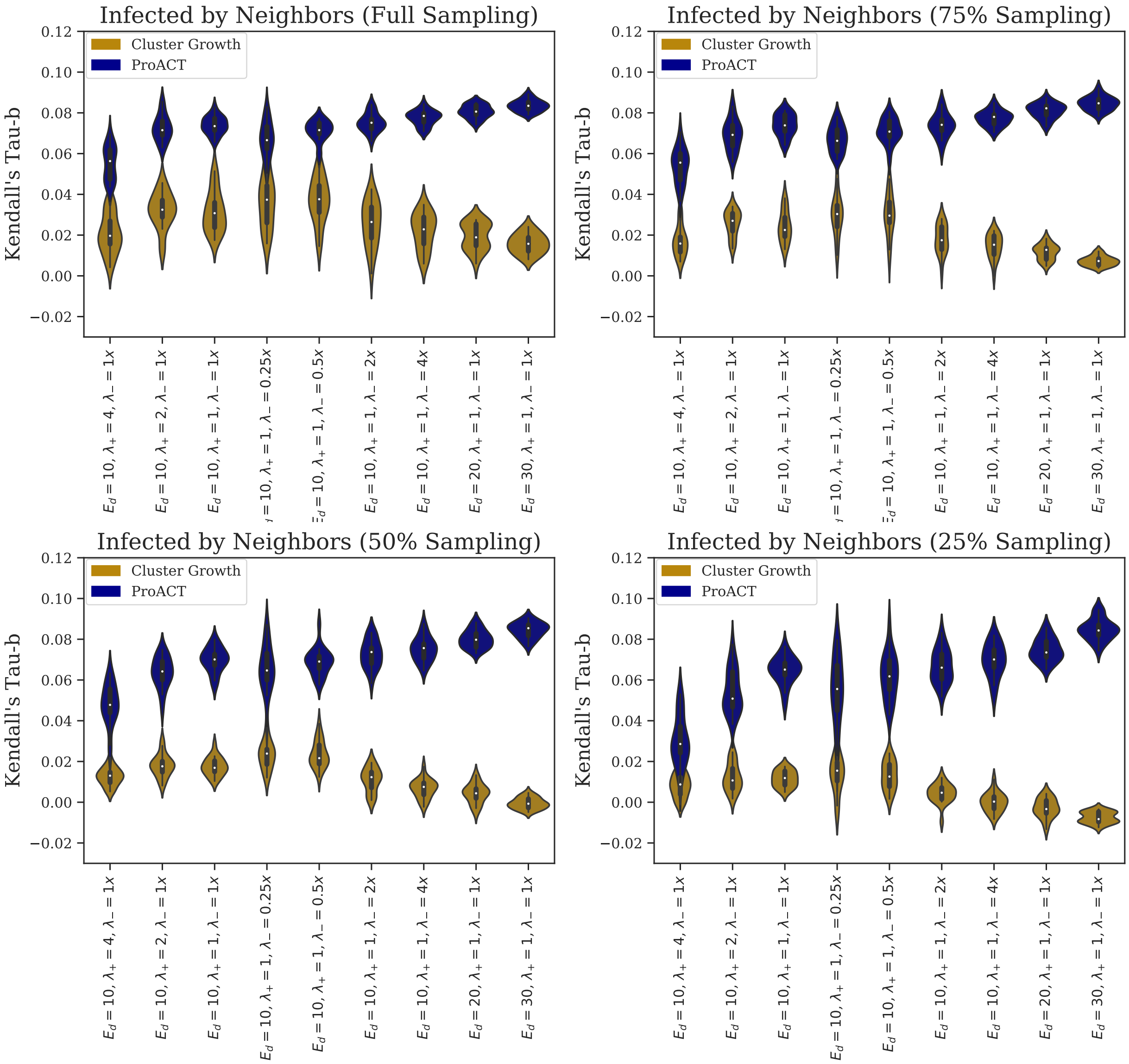


Fig. S8. Kendall Tau-b correlation between the optimal ordering of samples (i.e., based on their number of transmissions in year 10) and the orderings by the two prioritization methods, where effectiveness is measured by counting the number of future transmissions by all neighbors of a given individual. Distributions are across 20 replicates and are shown for each simulation condition. Results are shown for full sampling and subsampled data.

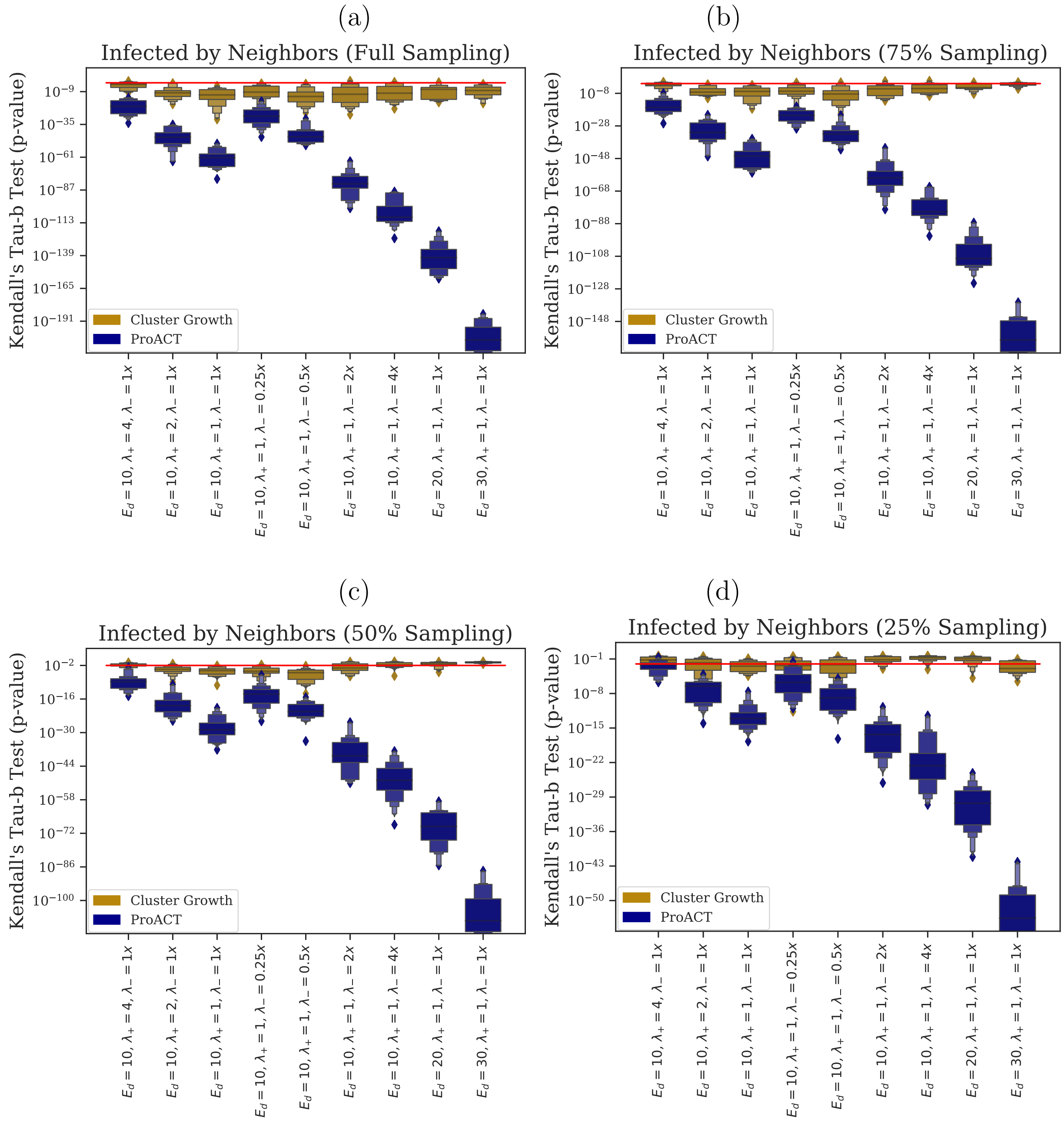


Fig. S9. $p$-values of one-sided Kendall Tau-b correlation tests between the optimal ordering of samples (i.e., based on their number of transmissions in year 10) and the orderings by the two prioritization methods, where effectiveness is measured by counting the number of future transmissions by all neighbors of a given individual. Distributions are across 20 replicates and are shown for each simulation condition. Results are shown for full sampling and subsampled data. See Figure S6 for Tau-b test statistic distributions.

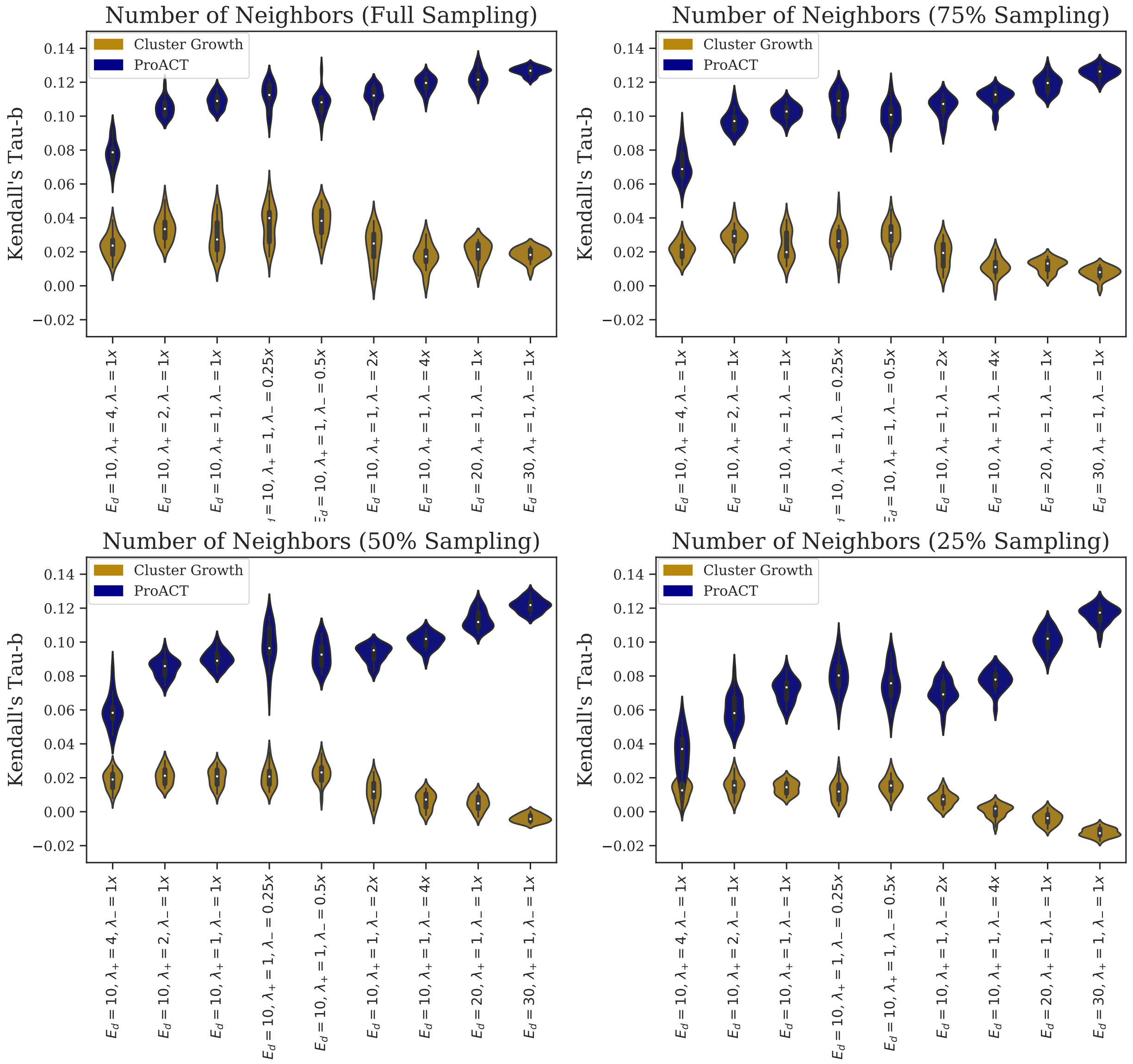


Fig. S10. Kendall Tau-b correlation between the number of social contacts samples and the orderings by the two prioritization methods, where effectiveness is measured by counting the number of future transmissions by all neighbors of a given individual. Distributions are across 20 replicates and are shown for each simulation condition. Results are shown for full sampling and subsampled data.

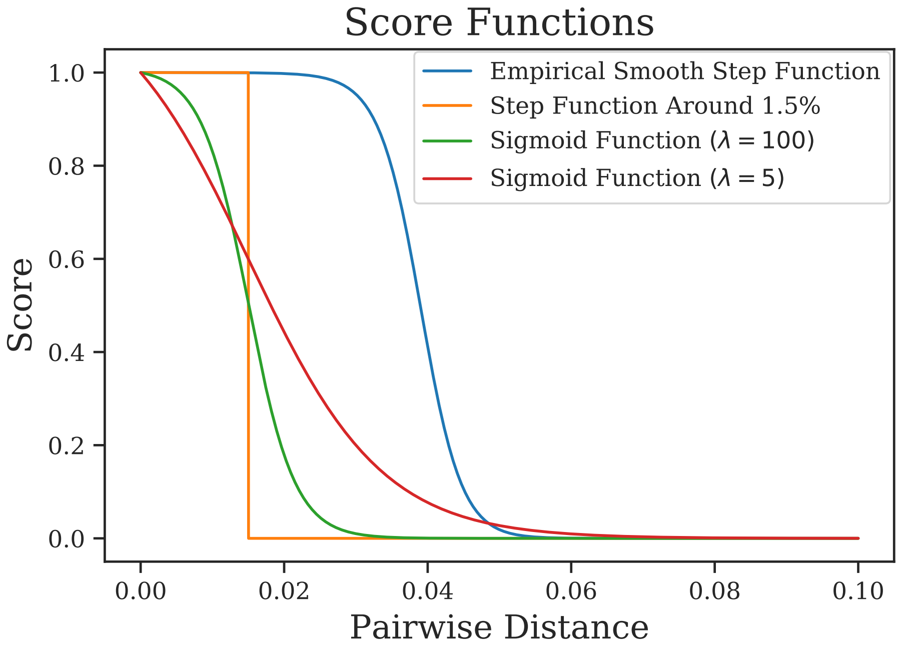


Fig. S11. Score functions vs. pairwise sequence distance.

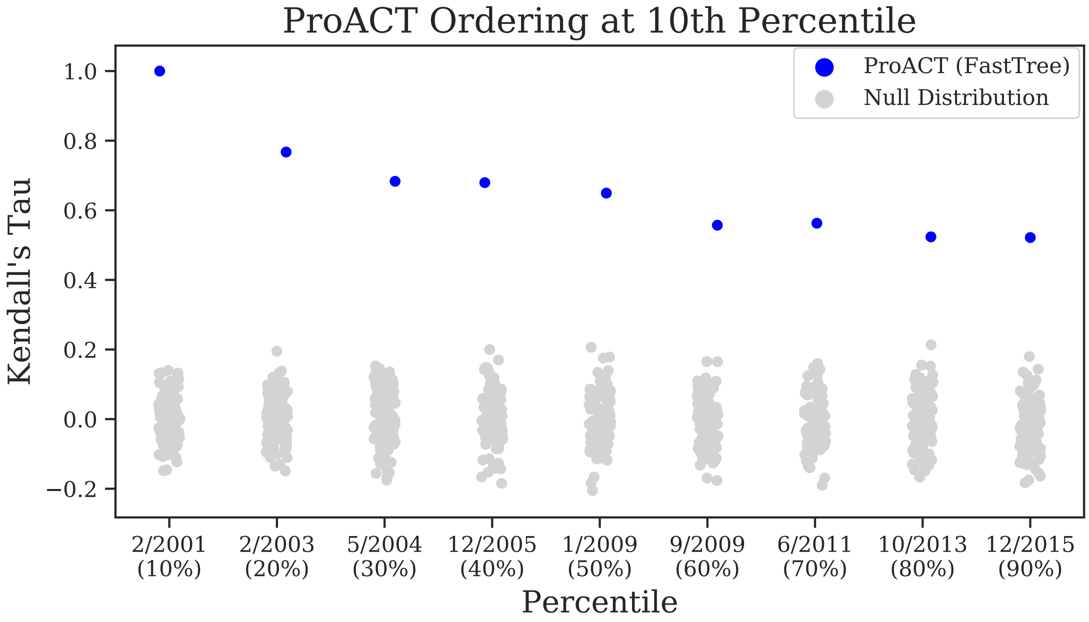


Fig. S12. Kendall's tau-b test results for ProACT ordering with respect to the ProACT ordering obtained with only the first decile of the datasat. The full San Diego dataset was split into two sets (*pre* and *post*) at each decile (shown on the horizontal axis). The individuals in *pre* were ordered using ProACT and by cluster growth. Kendall's tau-b correlation coefficient was computed for each ordering with respect to the ProACT ordering at the first decile. The null distribution was visualized by randomly shuffling the individuals in *pre*.

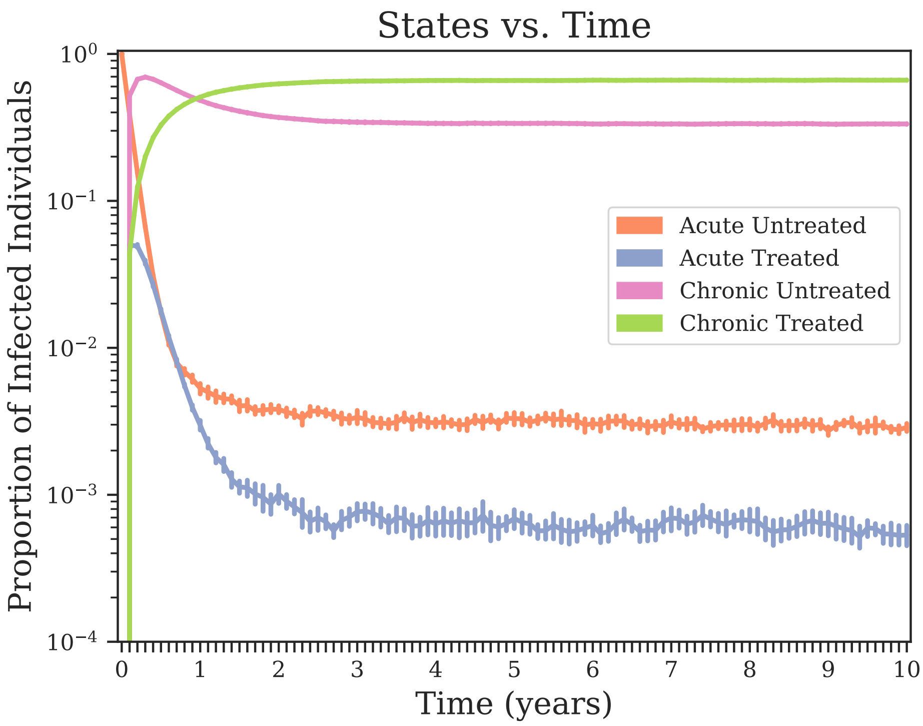


Fig. S13. Proportion of individuals in each infected state (AU, AT, CU, and CT) vs. time in simulations in which all seed individuals at time 0 were placed in state AU.

Table S1. Kendall's tau-b test for a null hypothesis that a given prioritization yields a total outcome measure no better than random. We show $p$-values for a real San Diego dataset for the first through ninth deciles. These $p$-values do not correct for multiple hypothesis testing. Tests that failed to reject the null hypothesis with (uncorrected) $\alpha = 0.001$ are marked with †.

| | Sigmoid Function ($\lambda = 5$) | | | | | | | | |
|---|---|---|---|---|---|---|---|---|---|
| | 10% | 20% | 30% | 40% | 50% | 60% | 70% | 80% | 90% |
| GD + Cluster Growth | $6 \times 10^{-4}$ | $^{\dagger}6 \times 10^{-3}$ | $3 \times 10^{-7}$ | $5 \times 10^{-5}$ | $8 \times 10^{-6}$ | $2 \times 10^{-7}$ | $8 \times 10^{-8}$ | $1 \times 10^{-6}$ | $1 \times 10^{-10}$ |
| ProACT (FastTree) | $1 \times 10^{-8}$ | $8 \times 10^{-5}$ | $2 \times 10^{-6}$ | $5 \times 10^{-8}$ | $1 \times 10^{-8}$ | $1 \times 10^{-11}$ | $1 \times 10^{-10}$ | $3 \times 10^{-11}$ | $1 \times 10^{-17}$ |

| | Sigmoid Function ($\lambda = 100$) | | | | | | | | |
|---|---|---|---|---|---|---|---|---|---|
| | 10% | 20% | 30% | 40% | 50% | 60% | 70% | 80% | 90% |
| GD + Cluster Growth | $1 \times 10^{-8}$ | $2 \times 10^{-11}$ | $6 \times 10^{-20}$ | $3 \times 10^{-24}$ | $2 \times 10^{-23}$ | $5 \times 10^{-17}$ | $3 \times 10^{-15}$ | $6 \times 10^{-11}$ | $4 \times 10^{-16}$ |
| ProACT (FastTree) | $2 \times 10^{-10}$ | $7 \times 10^{-9}$ | $3 \times 10^{-11}$ | $4 \times 10^{-18}$ | $9 \times 10^{-17}$ | $4 \times 10^{-20}$ | $7 \times 10^{-15}$ | $2 \times 10^{-12}$ | $1 \times 10^{-20}$ |

| Parameter | Default Value |
|---|---|
| Number of Contact Network Communities | 20 |
| Number of Individuals per Community | 5,000 |
| Mean Number of Edges Within Community | 10 |
| Mean Number of Edges Outside Community | 1 |
| Number of Seed Individuals | 15,000 |
| Seed Selection Model | Uniformly Random |
| Seed State Frequencies $\{AU, AT, CU, CT\}$ | $\{0.0033, 0.0006, 0.3396, 0.6565\}$ |
| Expected Transition Time AU→CU | 6 weeks |
| Expected Transition Time AT→CT | 12 weeks |
| Expected ART Initiation Time | 1 year |
| Expected ART Termination Time | 25 months |
| Rates of Infectiousness $\{AU, AT, CU, CT\}$ | $\{0.1125, 0.005625, 0.0225, 0.000\}$ |
| Seed Sequence Phylogenetic Model | Non-Homogeneous Yule Process |
| Seed Phylogeny Height | 25 years |
| Seed Phylogeny Speciation Rate Function | $\exp(-t^2) + 1$ |
| Mutation Rate Model | Truncated Normal |
| Mutation Rate Location | 0.0008 |
| Mutation Rate Scale | 0.0005 |
| Mutation Rate Minimum | 0 |
| Mutation Rate Maximum | $\infty$ |
| Viral Sequence Type | HIV-1 Subtype B *pol* |
| Sequence Evolution Model | GTR+Γ |
| GTR State Frequencies $\{p_A, p_C, p_G, p_T\}$ | $\{0.392, 0.165, 0.212, 0.232\}$ |
| GTR Transition Rates $\{\lambda_{AC}, \lambda_{AG}, \lambda_{AT}, \lambda_{CG}, \lambda_{CT}, \lambda_{GT}\}$ | $\{1.766, 9.588, 0.692, 0.863, 10.283, 1.000\}$ |
| GTR Gamma Distribution Shape | 0.405 |
| Viral Population Growth Rate Model | Logistic |
| Viral Population Growth Rate | 2.851904 |
| Initial Viral Population Size | 1 |
| Viral Population T50 | -2 |
| Number of Sampled Lineages per Person | 1 |
| Time of Sampling | ART Initiation |

Table S2. Default FAVITES simulation parameters.
